# Practical Bayesian Inference in Neuroscience: Or How I Learned To Stop Worrying and Embrace the Distribution

**DOI:** 10.1101/2023.11.19.567743

**Authors:** Brandon S Coventry, Edward L Bartlett

**Affiliations:** Department of Neurological Surgery and the Wisconsin Institute for Translational Neuroengineering, University of Wisconsin-Madison, Madison, WI USA 53705; Weldon School of Biomedical Engineering, Department of Biological Sciences, and the Institute for Integrative Neuroscience, Purdue University, West Lafayette, IN USA 47907

**Keywords:** Bayesian Inference, Neural Data Analysis, Statistical Inference

## Abstract

Typical statistical practices in the biological sciences have been increasingly called into question due to difficulties in replication of an increasing number of studies, many of which are confounded by the relative difficulty of null significance hypothesis testing designs and interpretation of p-values. Bayesian inference, representing a fundamentally different approach to hypothesis testing, is receiving renewed interest as a potential alternative or complement to traditional null significance hypothesis testing due to its ease of interpretation and explicit declarations of prior assumptions. Bayesian models are more mathematically complex than equivalent frequentist approaches, which have historically limited applications to simplified analysis cases. However, the advent of probability distribution sampling tools with exponential increases in computational power now allows for quick and robust inference under any distribution of data. Here we present a practical tutorial on the use of Bayesian inference in the context of neuroscientific studies. We first start with an intuitive discussion of Bayes’ rule and inference followed by the formulation of Bayesian-based regression and ANOVA models using data from a variety of neuroscientific studies. We show how Bayesian inference leads to easily interpretable analysis of data while providing an open-source toolbox to facilitate the use of Bayesian tools.

**Significance Statement:** Bayesian inference has received renewed interest as an alternative to null-significance hypothesis testing for its interpretability, ability to incorporate prior knowledge into current inference, and robust model comparison paradigms. Despite this renewed interest, discussions of Bayesian inference are often obfuscated by undue mathematical complexity and misunderstandings underlying the Bayesian inference process. In this article, we aim to empower neuroscientists to adopt Bayesian statistical inference by providing a practical methodological walkthrough using single and multi-unit recordings from the rodent auditory circuit accompanied by a well-documented and user-friendly toolkit containing regression and ANOVA statistical models commonly encountered in neuroscience.

## Introduction

The scientific process generates new experiments and updates models based on quantitative differences among experimental variables, evaluated by statistical inference. Inference tools are foundational to these studies, providing the necessary machinery to make decisions and conclusions from data. Frequentist-based null significance hypothesis testing (NHST) has been the gold standard of inference in neuroscience and science at large in part due to the computational simplicity of frequentist models compared to permutation sampling or Bayesian-based methods. A significant problem present in the current practice of NHST, however, arises in the adoption of the p-value as the *de facto* metric of experimental “success”, notorious for its difficulty in interpretation and correct usage (Krueger and Heck, 2019). The confluence of exponential increases in computational power with the wider discussion of problems with NHST usage has created renewed interest in Bayesian inference as an alternative to frequentist NHST while offering interpretability benefits over the p-value and NHST overall.

The use of p-values thresholds as a ubiquitous decision rule in frequentist methods is fraught with problems due to fundamental misunderstandings of its use, interpretability, and most pathologically, its susceptibility to intentional and unintentional p-hacking(Nuzzo, 2014). Contrary to the initial intent of Ronald Fisher(Fisher, 1992), the p-value has often become the gatekeeper of significance in studies. In this role, it limits deeper observations into data, and it is often used without proper experimental design to ensure proper use and control. Statistical inference methods require first defining a statistical model with the power to adequately describe the data-generating process. Inference is then performed to estimate the population distribution from limited samples of observed data. Once estimates of population distributions are made, the determination of whether or not these distributions represent a significant effect is determined. NHST is somewhat a victim of its own success, where common practice has distilled the practice of NHST to chase the somewhat arbitrary p<0.05 measure of significance devoid of model or data considerations(Krueger and Heck, 2019). Furthermore, even in the best of experimental designs, the p-value is a surrogate for arguably what a researcher is most interested in: how likely is it that observed data has some effect different from null(Kruschke, 2011; Gelman and Shalizi, 2013).

Bayesian methods offer a solution to the problem of pathological p-value use and interpretation, providing a direct measure of the probability that observations have some effect(Kruschke, 2011; Gelman and Shalizi, 2013). This is done by reallocation of probability of possibilities as parameters in a mathematical model of the data-generating process, leading to probabilistic estimates desired by but not attainable with p-value analyses. Bayesian methods are inherently data-driven; models are built with prior knowledge directly incorporated from parameters estimated directly from observed data.

Bayesian inference, though chronologically younger than frequentist approaches, was not adopted as the primary inference paradigm due to the computational demands necessary to solve inference problems outside of certain canonical forms(Bishop, 2006) and the adoption of frequentist interpretation of probability(Fienberg, 2006). Inference on arbitrary distributions required a deeper mathematical knowledge and computation of integrals which were potentially intractable without modern numerical integration techniques. Frequentist paradigms however were more easily adapted to computationally simple algorithms, allowing researchers to “do statistics” without extensive formal training. However, exponential increases in computational power with the development of powerful Markov chain Monte Carlo (MCMC) sampling methods now allow researchers to perform meaningful Bayesian inference on arbitrary distributions underlying observed data(Gilks et al., 1996). While Bayesian approaches have received some attention for inference in electrophysiology(Wood et al., 2004; Wu et al., 2006; Vogelstein et al., 2009; Cronin et al., 2010; Gerwinn et al., 2010; Park and Pillow, 2011), the advantageous interpretability of inference and data driven nature found in Bayesian statistics have of as yet been widely used in neuroscientific studies. This is in part a pedagogical problem, in that most neuroscientists do not encounter Bayesian statistics during formal training combined with the perception that a high level of mathematical acuity is necessary to perform Bayesian inference.

The goal of this tutorial is to remedy the opacity that often accompanies discussions of Bayesian inference by providing simple, step-by-step walkthroughs of Bayesian inference with four common inference paradigms utilizing tools that facilitate Bayesian computation for users at all levels of mathematical skill. We also aim to demonstrate the explanatory power of Bayesian inference in the context of neuroscience data. While the aim of this article is focused on application, this tutorial will begin with a brief introduction to Bayes’ rule and its constituent components necessary for inference. For more theoretical and mathematical considerations of Bayesian inference, see the following books and articles(Gerwinn et al., 2010; Colombo and Seriès, 2012; Bielza and Larranaga, 2014; Kruschke, 2014; Kruschke and Vanpaemel, 2015; Ma, 2019; Gelman et al., 2021; Van De Schoot et al., 2021; Cinotti and Humphries, 2022).

### Outline of Bayesian Methods

To best facilitate practical application of Bayesian inference methods in neuroscience, a variety of datasets were acquired and analyzed. Data acquisition paradigms are described in materials and methods below. Bayesian inference is introduced in the context of regression to infer changes in inferior colliculus single-unit firing from changing auditory stimuli. T-test-like group comparisons are demonstrated using computational models of basal ganglia thalamocortical local field potentials. Bayesian implementations of repeated measures and random effects models are demonstrated using chronic, multi-channel single-unit recordings from auditory cortex. Finally, Bayesian ANOVAs and ANCOVAs are utilized in assessing age-related changes in inferior colliculus single unit firing. All model implementations are available in our Bayesian inference for neuroscience toolbox at https://github.com/bscoventry/Practical-Bayesian-Inference-in-Neuroscience-Or-How-I-Learned-To-Stop-Worrying-and-Embrace-the-Dist.

### An Introduction to Bayesian Inference

Bayes’ rule forms the basis for all Bayesian inference as the machinery to reallocate probability from prior distributions and observed data to the posterior. While a full mathematical and computational treatment of Bayes’ rule is out of scope for this article, we begin by outlining the components of Baye’s rule, how inference is performed, and present a hypothetical experiment to show how priors might be formed.

### Bayes’ Rule

Foundational to Bayesian approaches is a complementary, but epistemically differing view of probability from that of frequentist approaches. While the frequentist perspective treats probability as the relative frequency of the occurrence of some event, the Bayesian perspective instead treats probability as the expectation of an event occurring which can be used to not only quantify the state of knowledge of an event, but also the uncertainty involved in measuring an event(Bishop, 2006; Kruschke, 2011, 2014; Van De Schoot et al., 2021). Traditionally, the Bayesian perspective has been called ‘belief’(Kruschke, 2010), a perhaps unfortunate name which belies the fact that the Bayesian perspective of uncertainty of an event is fundamentally quantifiable. Perhaps a better description of Bayesian belief is instead quantification of the state of knowledge by accounting for uncertainty. The cornerstone of Bayesian inference is Bayes rule, defined as:

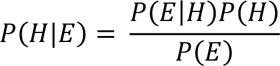

where H is the quantification of the state of a hypothesis, and E is the quantification of observed evidence. In the context of inference, it is helpful to explicitly state the role of the model in Bayesian formulations:

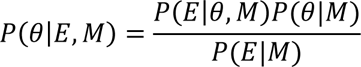

where M is the model of the data generating process and θ are the model parameters. The individual components of Bayes’ rule are given names corresponding to the purpose they serve, with *P*(θ|*E*, *M*) called the posterior distribution, *P*(*E*|θ, *M*) the likelihood function, *P*(θ|*M*) the prior distribution, and *P*(*E*|*M*) the evidence or marginal likelihood function. Taken together, Bayes’ equation represents the quantification of observed data accounting for prior knowledge(Fig 1A). Each component plays a key role in Bayesian inference and each will be discussed briefly below.

**Figure 1:**
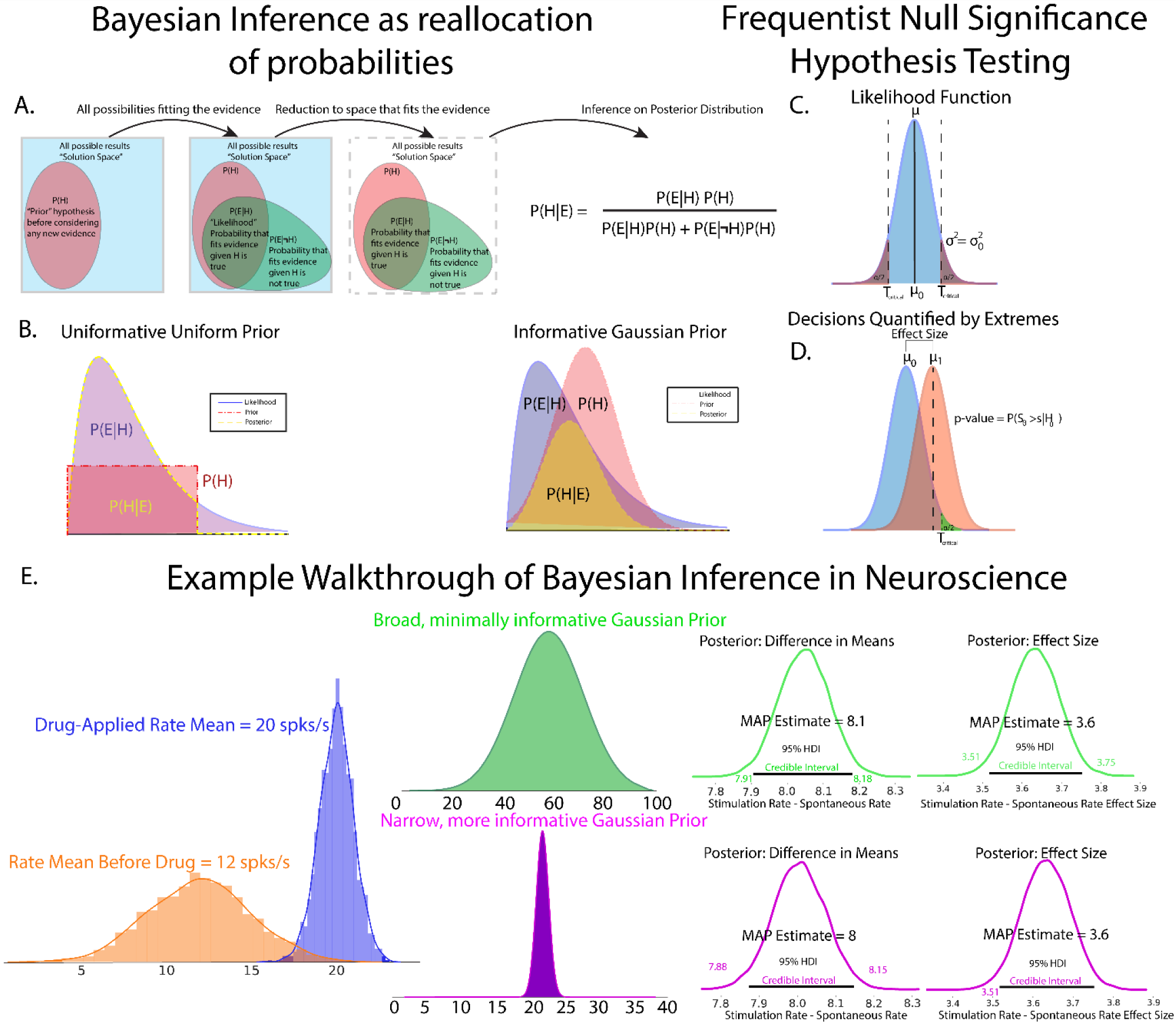
Graphical description of Bayes rule and the interaction between prior distributions and likelihood functions leading to the final posterior distribution. A. Bayes rule can be thought of as a reallocation of probability to the posterior after accounting for prior distributions and observed evidence. B. An example of posterior generated from an inverse-Gamma distributed likelihood and a uniformly distributed prior. Uniform priors reflect the likelihood function, and thus the observed data with no redistribution probability, making uniform distributions uninformative priors. However, care must be taken in using uniform distributions as observed data outside of prior bounds is mapped to 0 probability. A second example of a posterior generated from an inverse-gamma distributed likelihood and a gaussian distributed prior shows how a prior can be considered informative (right). In this posterior, the prior “shapes” the posterior to a greater extent than a uniform prior. Well designed priors in neuroscientific data can thus shape posteriors away from responses which are not physiological and can be vetted in the statistical decision, data review, and post-publication review stages. Prior distributions with longer tails can handle extremes of observed data by mapping extreme events to low, but non-zero representation in the posterior. Example B represents extremes of prior choices, with minimally informative priors often chosen to let the data “speak for itself” with little change to posterior from prior influence. C. Comparison of Bayesian and Frequentist statistics. NHST utilizes a parameterized likelihood function that describes the data generating process. D. Decisions in NHST are generally made based on quantification of the probability of observing test results as extreme as observed data assuming that the null hypothesis is true. E. An illustrative example of how Bayesian inference might be used in an electrophysiological experiment. Prior distributions can be iteratively formulated by previous experiments drawn from both internal and published data.

### The Model Evidence

The denominator term *P*(*E*|*M*), called the model evidence (or just the evidence or marginal likelihood in Bayesian parlance) is the quantification of the probability of observing the data under a chosen model of the data generating function. At first glance, the calculation of the total evidence appears to be an insurmountable task. In reality this term is the weighted average of parameter values in a given model weighted by the relative probability of a given parameter value(Kruschke, 2014) and thus acts as a normalization term to ensure the numerator is a proper probability distribution. The structure of *P*(*E*|*M*) will change based on whether the distributions represent probability mass functions (discrete case) or probability density functions (continuous case). In the discrete case, the evidence is

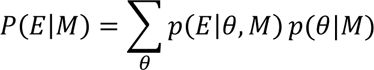

and in the continuous case:

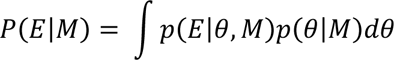

The evidence function thus represents an average of the likelihood function across all parameter values conditioned on the prior distribution. The marginal likelihood can also be utilized to assess the plausibility of two competing models(Johnson et al., 2023). The evidence, especially in the continuous case, is historically what made Bayesian inference difficult due to the need to evaluate a complex integral numerically. However, the advent of Markov-chain Monte Carlo (MCMC) methods with improvements in personal computer processing power has allowed for computationally efficient integration without the need for supercomputing hardware. MCMC methods will be discussed in a subsequent section.

### The Prior

The prior, *P*(θ|*M*), is often the major stumbling block for those entering into Bayesian inference, but this hurdle is less about the prior, and more about what the prior is perceived as. The prior, *P*(θ|*M*) describes the investigators prior beliefs on the state of knowledge of the study. While normally distributed priors are common, the prior can take the form of any proper probability distribution. Critics of Bayesian inference have described the prior as purely subjective, but we, and many others(Kruschke, 2010; Box and Tiao, 2011; Gelman and Shalizi, 2013), argue that the prior represents an explicit declaration of the investigators knowledge, assumptions, and the general state of a field which is implicit and often is present but not stated in frequentist approaches. Moreover, one is encouraged to perform prior predictive checks to compare the sensitivity of competing priors in a Bayesian inference model, as we will show subsequently. The practice of the design of experiments and their resulting publications are rife with implicit priors which are often not acknowledged or realized when reporting results. As an example, consider study of cortical extracellular single unit recordings(Paninski et al., 2004; Bartlett and Wang, 2007; De La Rocha et al., 2007; Coventry et al., 2024 as illustrative examples). The investigator could be leading a project with vast knowledge accumulated over years of study. Or the investigator is a trainee of a career researcher who draws a view of cortical physiology from their experienced mentor mixed with reading current literature. When designing an experiment, the investigator will have some intuition regarding likely and biologically feasible resting state and stimulus-evoked firing rates, cognitively assigning relatively low likelihood of seeing extremes of firing rates with higher likelihood assigned to moderate firing rates previously observed in literature or seen in experiments, and likely will discard or treat as outliers firing rates on the extremes or thought to be non-biological noise. The power of the prior distribution in Bayesian approaches is in part the need to explicitly quantify and report these prior beliefs, which can be analyzed and scrutinized as part of the peer review or post-publication process. Prior distributions also require investigators to consider their biases and relative expectation on the importance of previously recorded and read data, promoting a deeper understanding of not only the data obtained within their lab, but also of the general state of the specific neuroscience field. As the name implies, prior beliefs are quantified as probability distributions by the investigators.

This begs the question as to what a prior might look like in newer avenues of study where a paucity of data exists. Or in situations where researchers and data analysts want the data to “speak for itself” outside any influence of the prior. In these cases, priors can be designed to be “non-informative” or “weakly-informative”, assigning broad, non-committal distributions to the prior. One might assign a uniform distribution on the prior, effectively treating each parameter outcome as equally likely. Uniformly distributed priors do require some caution, however, as any parameter value outside of the bounds of the uniform distribution is automatically assigned probability 0 in the posterior, even if that value has been observed(Fig 1B Left). In many cases, it’s better to allow small, but nonzero probabilities to extreme values, such as the tails of a normal distribution, such that evidence for unexpected events is represented in the posterior given strong data(Fig 1B Right). Conversely, priors can be made to be highly informative in situations where physiological bounds are well known and well-studied, where extreme values are known to be biophysically irrelevant or impossible or known to be due to instrument noise(e.g. large 50/60 Hz noise peak in power spectrum indicative of wall power noise).

### The Likelihood

The likelihood function, *P*(*E*|θ, *M*) describes the probability that data is observed given parameter values θ in a data generating model M. In the context of inference, the likelihood function updates information given in a prior distribution to the posterior distribution given the observed data(Etz, 2018). The likelihood function is generally not a proper distribution, in that it is conditioned on yet unknown parameters and may not integrate to 1, but the evidence and prior terms ensures that resultant posterior distributions are true probability densities. The idea of likelihood functions are present in both Bayesian and frequentist models, but have vastly different interpretations. The model parameters in a frequentist viewpoint converge upon singular values learned, usually though maximum likelihood estimation, from merging competing hypotheses of data. Bayesian approaches treat model parameters as ranges arising from distributions after observing the data at-hand.

### The Posterior

The prior, likelihood, and evidence then form the posterior *P*(θ|*E*, *M*), the reallocation or mapping of probability from likelihood function, prior, and model evidence to an all-encompassing distribution. The posterior thus is the evidence for parameters θ conditioned on observed data and a model of the data generating function. The posterior forms the basis for inference, with all relevant information encoded in its distribution. Inference on the posterior distribution is covered in a section below.

### Estimation of the Posterior

Despite Bayes’ rule being formulated before Fisher’s description of frequentist methods, a major reason that Bayesian inference was not been widely adopted was fundamentally a computational one, in that evaluation of Bayes’ rule often requires solving non-trivial integrals. A subset of computationally tractable prior distributions and likelihood functions formed canonical posteriors in which the posterior is easily inferred. However, these cases are not generalizable to experimental data which can be noisy and not well behaved. Modern Markov-chain Monte-Carlo (MCMC) tools have been developed to quickly and easily estimate arbitrary distributions. MCMC involves the generation of random samples which converge to a target probability distribution, the details of which can be learned from the following reviews(Hoffman and Gelman, 2011; Betancourt, 2017).

### Making Decisions on the Posterior

We define inference broadly as the process by which reasoning and decisions about a phenomena are made from a sample of observations of the phenomena. Incorporation of prior knowledge in Bayesian inference allows for optimal decision making on observed data(Blackwell and Ramamoorthi, 1982). The posterior contains all necessary information to make inferences on experimental data incorporating prior knowledge. However, it is best to consider the specific goals of inference before performing statistics. Possible goals of inference are as follows(Kutner et al., 2005; Kruschke, 2014):

- Infer the parameters of a model.
- Reject or confirm a hypothesis, null or otherwise
- Compare two or more competing models

In the case of neuroscientific studies of unit electrophysiology, inferring model parameters occurs when an experiment aims to establish how neural firing rates change with changes in applied stimuli. Or one may want to confirm or reject a null hypothesis that a treatment has the desired effect or that there are differences between neural populations. Importantly, because the Bayesian inference operates solely on the posterior distribution, one can confirm or reject competing hypotheses and not simply reject the null as in frequentist NHST.

Regardless of the goal, inference always involves analyzing the posterior, which provides a complete representation of the distribution of a given parameter given the experimental data. Therefore, decisions about the data, the effect of model parameters, and/or which hypothesis has more evidence is performed with calculations on the posterior. There are multiple decision rules that can be used to assess the posterior. The most common, and in our opinion, the most intuitive is that of the Bayesian credible interval. The confidence interval calculates the probability that a population parameter lies in a certain interval. As credible intervals are not strictly unique, Bayesian inference convention is to fix the interval to the smallest interval which contains 95% of the posterior distribution density mass called the highest density interval (HDI). Observations of posterior HDIs can then be used to assess the relative effect of a parameter. Regions of practical equivalence (ROPE) may be included in the posterior distribution that explicitly define a range of values that are effectively equivalent to a null value, with parameters considered significant if 95% of the posterior distribution (95% HDI) does not contain 0 or any values in the ROPE (Kruschke, 2018). Along with posterior HDIs, calculations of maximum *a posteriori* (MAP, distribution mode) estimates from the posterior are performed to quantify a most likely parameter value. While decision rules are important to assess the relative effect of statistical model parameters, we reiterate that simply passing a decision rule should not conclude the inference step. Inference should be made in context of the evidence presented in model quality checks, observed data posterior distributions, and decision metrics.

### A Simple Example of Bayesian Applications in Neuroscience

As a simple, motivating example of how priors and likelihoods can be derived in neuroscientific studies, consider a hypothetical *in vitro* whole-cell patch clamp experiment recording a cortical neuron (Fig 1E). The experimental details are less important than the formulation of Bayesian models, but are given in some detail to help visualize the hypothetical study. Consider a study to assess the effect of a novel drug on firing rates of a specific class of cortical neuron. The goal of the study is to assess the change in the firing frequency versus injection current before and after drug application during current-clamp recording. Before the beginning of the experiment, the graduate student surveys the knowledge (i.e. a literature search) surrounding these neurons and assigns a broad, normally distributed prior centered around the average firing rates of these cortical neurons across all studies surveyed. The logic to this prior is that while firing rate distribution to this novel drug is not known, the totality of observed firing rates reasonably approximates the range of possible firing states of that neuron. The graduate student runs the study, plots the distribution of observed results and finds that drug application produces small increases in mean firing rates that are very consistent between trials and cells, as evidenced by relatively tight distributions of data (Fig 1E left, observed data distributions). However, the overlap of naive and drug conditions means that quantification of the uncertainty of measurement of the drug effect must be performed in the form of the MAP and credible intervals in each condition. The distributions of evoked firing rates as a function of injected currents are the data encoded in the likelihood function. After completion of the experiment, the advisor of the graduate student noticed that both *in vitro* and *in vivo* data were included in the prior, even though firing rates differ dramatically between those preparations in previous work. Re-running the analysis with the highly informed prior based on in vitro data produces a similar result, but with a slight reduction in the uncertainty in the difference of group means as measured by group mean difference MAP estimates and credible intervals. In this case, comparing weakly and strongly informed models also shows that relatively drastic changes in the prior make minor changes in posterior estimates, suggesting that observed data is the primary driver of inference and is not unduly influenced by the prior. Taken together, both broad, minimally informative and highly informative priors often lead to similar decisions on the data and are useful for cases where prior knowledge is sparse. However, when prior knowledge is available, incorporating more information in the form of an informative prior backed by data provides reduction in uncertainty in estimation of study effects.

This simple example shows one way in which priors and likelihoods are defined in neuroscience studies in a manner that most researchers unknowingly do by running experiments and reading the literature. The prior serves to take intrinsic prior knowledge and quantify it to provide more transparent reporting of experimental results. Furthermore, data forms the likelihood function, facilitating inference that makes decisions on the data at hand while accounting for prior knowledge of the state of a particular field. It also introduces the idea of using model comparison to assess prior influence on inference, which will be rigorously explored below.

### Inference Using Null Significance Hypothesis Testing

While theoretical comparisons of Bayesian inference vs NHST are out of scope for this article, a brief description of decision-making using NHST is warranted to orient Bayesian inference. For detailed descriptions and discussions of NHST, see the following reference texts and articles(Kutner et al., 2005; Nieuwenhuis et al., 2011; Maris, 2012; Kruschke, 2014; Bzdok and Ioannidis, 2019). Proper NHST experimental design is performed before data acquisition with sample size, statistical tests, and significance levels set to achieve a desired level of statistical power. However, proper NHST experimental design is largely not followed, in part due to lack of proper statistical training and external career pressure(Munafò et al., 2017) as well as the relative difficulty in data acquisition(Button et al., 2013). We believe that Bayesian approaches provide a way to do statistics more in line with current neuroscientific practice, in which beliefs about data are updated as data is acquired, informed by the current state of the field. Statistical power relies on estimates of population variances and differences in group means. We note that this is somewhat analogous to the declaration of prior distributions in Bayesian inference. After data is acquired using the predefined stopping rule, NHST inference is then performed. Similar to Bayesian inference, NHST utilizes a parameterized likelihood function(Fig 1C) that describes the data-generating process. The null hypothesis is the likelihood function descriptive of a nonexistent effect. Statistical significance is most commonly assessed through the p-value metric, a measure quantifying the probability of obtaining a value of a given test statistic at least as extreme as the test statistic observed under the assumption of a null hypothesis. Another common metric for statistical significance and parameter estimation is the confidence interval, which utilizes sample distributions to estimate parameters of the population distribution. Importantly, unlike data-driven Bayesian approaches, both p-values and confidence intervals draw inference using in part data that is not observed, owing from calculations over hypothetical replications of an experiment(Wagenmakers, 2007; Kruschke, 2014). Comparisons between Bayesian methods and NHST are summarized in Table 1.

**Table 1:**
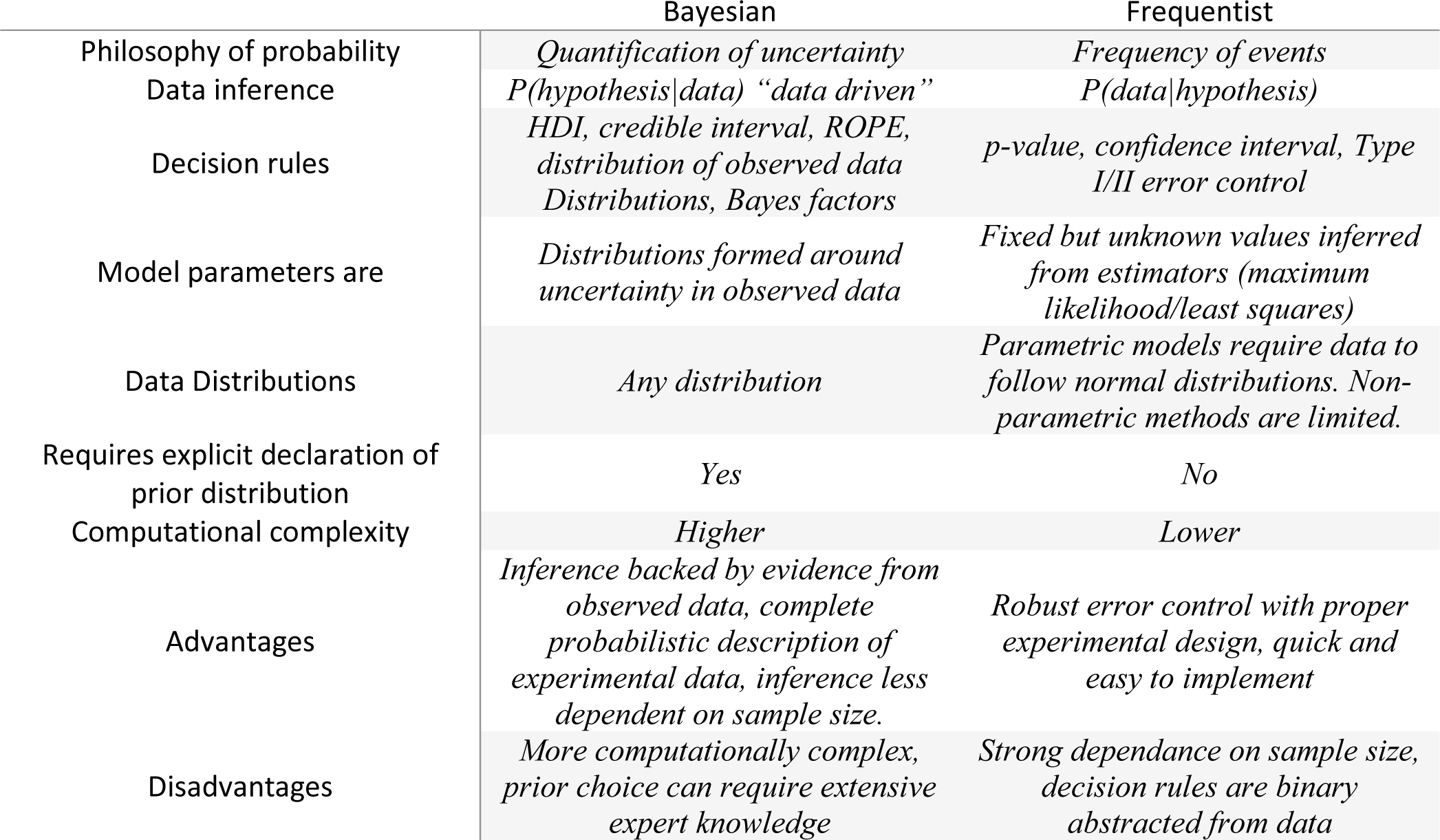
Brief Comparison of Bayesian and frequentist inference paradigms.

### Error Quantification and Model Comparison

Critical to any statistical model and inference therein is its fit to observed data. While it is entirely possible to perform linear regression on data distributions which are highly nonlinear, predictions and inference made by the model will likely be inaccurate. Both Bayesian and frequentist inference offer robust model error quantification. Bayesian approaches, however, can utilize the posterior distribution to not only quantify and bound the distribution of model errors, but also include *post hoc* posterior predictive sampling as part of the inference paradigm. Posterior predictive sampling involves making random draws from the posterior and building a sampling distribution. This distribution is then compared to the observed data distribution to quantify the model’s disparity from observed data. Along with posterior predictive checks, prior predictive checks act as a sensitivity measure of the influence of the prior distribution on the posterior distribution. Taken together, Bayesian inference thus allows for robust statistical inference on observed experimental data which appropriately includes prior knowledge of the state of the field.

### Formulation of Models and Applied Bayesian Inference

There are a multiplicity of programs and programming languages that facilitate Bayesian analysis, such as standalone programs of Jasp(Love et al., 2019) and probabilistic programming language packages such as BUGS(Brooks, 2003) and STAN(Carpenter et al., 2017), we chose to use PyMC(Salvatier et al., 2016) for its ease in explicitly declaring probability distributions and its implementation in Python which is in common use in neuroscientific data analysis. Model formation is often conserved between frequentist and Bayesian approaches; it is only the mode of inference that differs. However, for clarity, we will discuss both model formation and performing inference in the subsequent sections.

## Materials and Methods

Bayesian inference was performed on a range of data typical to neuroscience experiments. Regression models, ANOVA models, and group comparisons are performed on single-unit activity recorded from inferior colliculus (IC) neurons in response to auditory stimuli in young and aged rats(Palombi et al., 2001; Simon et al., 2004; Rabang et al., 2012; Herrmann et al., 2017). Random-effects regression models are performed on single units recorded in the auditory cortex (A1) using high-density recording arrays in response to infrared neural stimulation(Izzo et al., 2007; Cayce et al., 2011, 2014; Coventry et al., 2024) of the medial geniculate body (MGB). All surgical procedures used in this study were approved by the Institutional Animal Care and Use Committee (PACUC) of Purdue University (West Lafayette, IN, #120400631) and in accordance with the guidelines of the American Association for Laboratory Animal Science(AALAS) and the National Institutes of Health guidelines for animal research

### Computational Hardware

To underscore that meaningful Bayesian inference does not require cluster computing or extensive computational resources, all computations were performed on a Windows MSI GS-66 laptop with an Intel i7 processor with an Nvidia RTX2070 GPU. Our inference programs are CPU-bound, not requiring any GPU resources. Computations can be performed on most modern CPUs, but are accelerated with more CPU threads and cores and parallelization on GPUs.

### Bayesian Linear Regression and Analysis of Covariance Assessment of the Disruption of Temporal Processing in the Inferior Colliculus Due to Aging

The inferior colliculus (IC) is the major integrative center of the auditory pathway, receiving excitatory inputs from ventral and dorsal cochlear nuclei, excitatory and inhibitory inputs from the lateral and medial superior olivary complex(Kelly and Caspary, 2005) and inhibitory inputs from superior paraolivary nucleus and the dorsal and ventral nuclei of the lateral lemniscus(Cant and Benson, 2006; Loftus et al., 2010). The IC encodes auditory information through hierarchical processing of input synaptics with local IC circuitry(Caspary et al., 2002; Rabang et al., 2012; Grimsley et al., 2013). Age-related changes in auditory processing primarily arise as deficits in temporal processing(Frisina and Frisina, 1997; Parthasarathy et al., 2010; Parthasarathy and Bartlett, 2012; Herrmann et al., 2017). This dataset is composed of single unit responses recorded from young (Age≤ 6 months) and aged (age ≥ 22 months) Fisher 344 rats. Auditory brainstem responses were recorded from animal subjects a few days prior to surgery to ensure hearing thresholds were typical of the rodent’s age. Single unit recordings were performed in a 9’x9’ double-walled, electrically isolated anechoic chamber (Industrial Acoustics Corporation). Animals were initially anesthetized via a bolus injection of ketamine (VetaKet, 60-80 mg/kg) and medetomidine (0.1-0.2 mg/kg) mixture via intramuscular injection. Oxygen was maintained via a manifold and pulse rate and blood oxygenation monitored through pulse oximetry. Supplemental doses of ketamine/medetomidine (20 mg/kg ketamine, 0.05 mg/kg medetomidine) were administered intramuscularly as required to maintain surgical plane of anesthesia. An incision was made down midline and the skull exposed. Periosteum was resected and a stainless steel headpost was secured anterior to bregma via 3 stainless steel bone screws. A craniectomy was made above inferior colliculus (-8.5 anterior/posterior, 1 mm medial/lateral from bregma). A single tungsten electrode was advanced dorsally towards the central nucleus of the inferior colliculus (ICC) during which bandpass noise (200 ms, center frequencies 1-36kHz in five steps per octave, 0.5 octave bandwidth) was delivered. ICC was identified based on short-latency driven responses to bandpass noise search stimuli with ascending tonotopy and narrowly tuned responses to pure tones of varying frequencies. Once neurons were identified, responses from 5-10 repetitions of sinusoidal amplitude-modulated tones (750 ms tone length, modulation depth between -30 to 0 dB) were recorded using a preamplifying headstage (RA4PA, Tucker-Davis Technologies) and discretized at a sampling rate of 24.41 kHz (RZ-5, TDT). Sinusoidal amplitude-modulated tones were defined as:

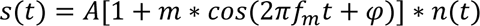

where m is modulation depth ranging between 0.032 − 1 (−30 – 0 *dB*, *f*_*m*_ the modulation frequency, φ the reference phase of the modulator, A is the scaling factor for stimulus sound level, and *n*(*t*) the broadband noise stimulus. SAM stimuli are schematized in Fig 2C. Single units were filtered between 0.3 and 5 kHz. Offline spike sorting was performed using OpenExplorer (TDT).

**Figure 2:**
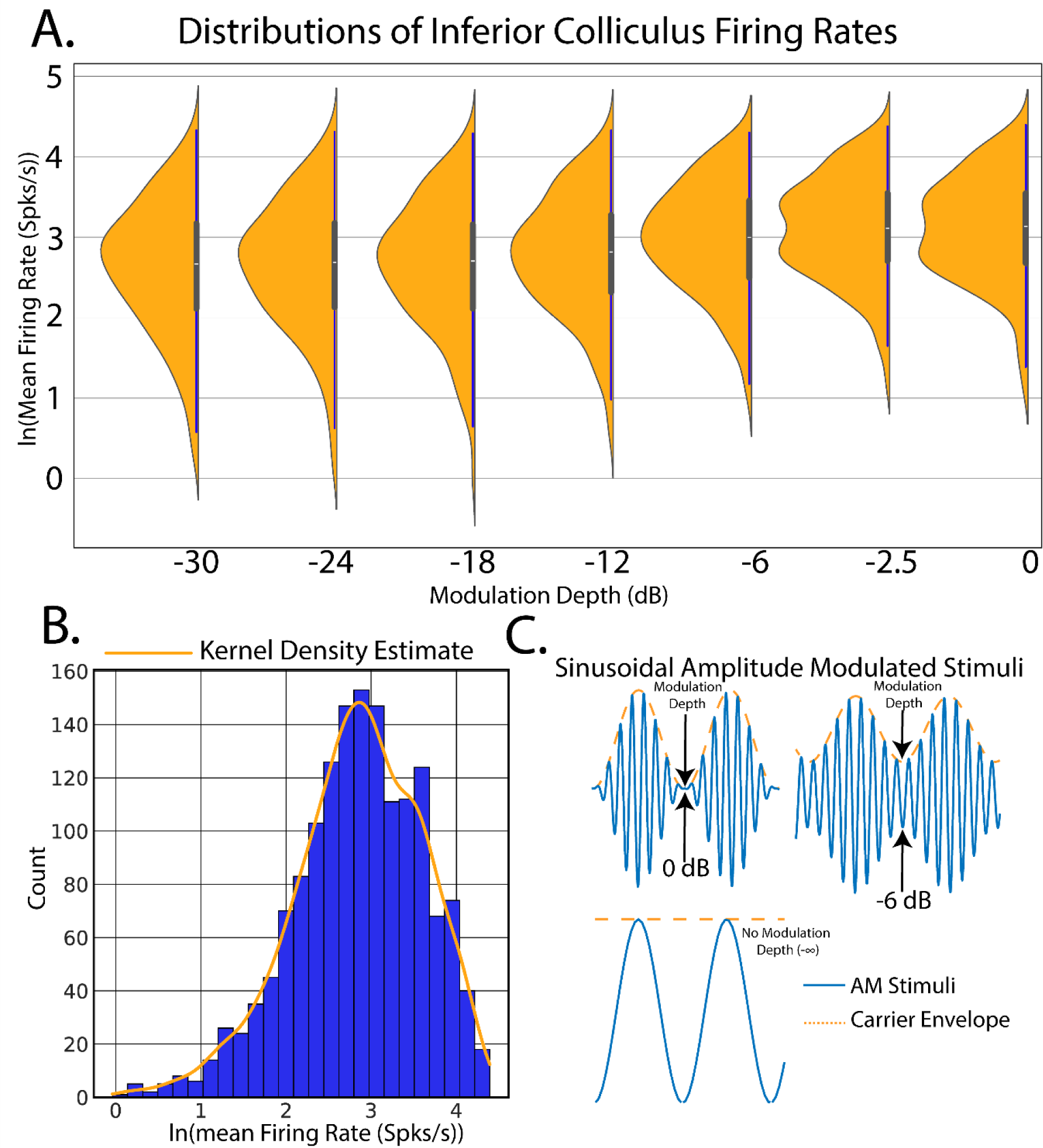
Example of Bayesian simple linear regression on population estimates of firing rate vs amplitude modulation depth stimuli. This model was applied to population single unit firing rates elicited from inferior colliculus with sinusoidal amplitude modulated (SAM) tones. The goal of this model was to predict evoked firing rates from increases in SAM modulation depths. A. Density plot of observed firing rates vs SAM modulation depth and fitted regression estimates. B. Histogram and kernel density estimation (KDE) of the distribution of SAM-evoked inferior colliculus firing rates. C. Schematic of amplitude modulated stimuli.

### Bayesian Multilinear Regression Assessment of Thalamocortical Recruitment From Infrared Neural Stimulation

Infrared neural stimulation (INS) is an optical technique using coherent infrared light to stimulate nerves and neurons without the need for genetic modification of the target or direct contact with tissue that offers spatially constrained activation above electrical stimulation(Wells et al., 2005, 2005; Izzo et al., 2007; Cayce et al., 2011, 2014; Coventry et al., 2020, 2024). In this study, rats were chronically implanted in A1 with 16 channel planar Utah-style arrays (TDT, Alacua FL) and stimulating optrodes in the medial geniculate body of auditory thalamus (Thor Labs, Newton NJ). Rodents were initially anesthetized with a bolus injection of a ketamine (80 mg/kg) and medetomidine (0.2 mg/kg) cocktail. Oxygen was maintained via a manifold and pulse rate and blood oxygenation monitored through pulse oximetry. Supplemental doses of ketamine/medetomidine (20 mg/kg ketamine, 0.05 mg/kg medetomidine) were administered intramuscularly as required to maintain surgical plane of anesthesia. An incision was made down midline and the skull exposed. The periosteum was removed via blunt dissection and 3 stainless steel bone screws were placed in skull for headcap stability. An additional titanium bones crew was placed in skull to serve as a chronic ground and reference point for recording electrodes. Craniectomies were made above medial geniculate body (-6 anterior/posterior, -3.5 medial/lateral from bregma) and auditory cortex (-6 anterior/posterior, -5 medial/lateral from bregma). Fiber optic stimulating optrodes were placed in the midpoint of MGB (-6 dorsal/ventral from dura) and affixed to the skull using UV-curable dental acrylic (MidWest Dental). A 16 recording channel planar array was putatively placed in layers 3/4 of auditory cortex, with placement confirmed by short-latency high amplitude multiunit activity elicited from band pass noise (200 ms, center frequencies 1-36kHz in five steps per octave, 0.5 octave bandwidth) test stimuli. Recording electrodes were sealed onto the headcap. Animals were allowed to recover for 72 hours prior to the beginning of the recording regime. All recordings were performed in a 9’x9’ electrically isolated anechoic chamber. During recording periods, animals received a intramuscular injection of medetomidine(0.2 mg/kg) for sedation. Optical stimuli were delivered from a 1907 nm diode laser (INSight open source optical stimulation system) coupled to the optrode with a 200 μm, 0.22 NA fiber (Thor Labs FG200LCC). Laser stimuli were controlled via a RX-7 stimulator (TDT) and consisted of train stimuli with pulse widths between 0.2-10 ms, interstimulus intervals between 0.2-100 ms and energy per pulse between 0-4 mJ. Applied laser energies were randomized to limit effects from neural adaptation with 30-60 repetitions per pulse width/interstimulus interval combinations. Signals from recording electrodes were amplified via a Medusa 32 channel preamplifier and discretized and sampled at 24.414 kHz with a RZ-2 biosignal processor and visualized using Open-Ex software (TDT). Action potentials were extracted from raw waveforms via real-time digital band-pass filtering with cutoff frequencies of 300-5000 Hz. Single units were extracted offline via superparamagnetic clustering in WaveClus (Quiroga et al., 2004). Studies were performed to assess the dose-response profiles of optically-based deep brain stimulation over the span of several months. As each electrode recorded diverse populations of neurons which are potentially subject to change due to electrode healing in, age of the device, and adaptation to the stimulus, a within subjects, repeated measures regression model was warranted. Bayesian hierarchical regressions can easily deal with complex models such as these. This data was part of a previous study(Coventry et al., 2024).

### Bayesian T-Test Assessment of Computational Models of Basal Ganglia Thalamocortical Function in Parkinson’s Disease

Parkinson’s disease is a chronic and progressive neurological disorder resulting from a loss of dopaminergic neurons in the substantia nigra of the basal ganglia circuit(Dickson, 2018; Bove and Travagli, 2019). Computational models of basal ganglia thalamocortical function provide insight into alterations of circuit dynamics as well as potential therapeutic targets. In prior work, a modified mean-field basal ganglia thalamocortical (BGTC) model (Grado et al., 2018; Coventry and Bartlett, 2023) was implemented to study network deep brain stimulation encoding mechanisms in dopamine depleted states. In the case of BGTC circuits, mean-field modeling consists of the average extracellular electric field response of collections of neurons with cellular properties modeled after each stage of the BGTC circuit in both healthy and dopamine depleted states(Van Albada and Robinson, 2009). In each model trial, local field potentials (LFP) were recorded from the globus pallidus internus in resting state and stimulation conditions. Stimulation trials consisted of subthreshold (0.01 mA, 100μs pulse width, 130 Hz) or suprathreshold (1.8 mA, 240μs pulse width, 130 Hz) deep brain stimulation of the subthalamic nucleus. LFP activity in the β band (13-30 Hz) is a known biomarker for Parkinson’s symptomology(Little et al., 2013; Guidetti et al., 2021), so LFP power spectral density estimates of β band activity were calculated. Total LFP power in the β band was calculated as follows:

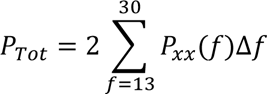

where *P*_*xx*_ is the power spectral density at frequency f, Δ*f* reflecting model sampling rate, and the factor 2 accounting for frequency folding from Fourier decomposition.

## Results

### Estimation of Spike Rates from Auditory Stimuli: A Motivating Example

To facilitate the discussion of Bayesian inference in neuroscience, consider an example found prominently in auditory neuroscience(Fig 2). In our first experiment, single unit recordings were made from the inferior colliculus (IC) in response to applied sinusoidal amplitude-modulated tones (SAM, see Methods). The goal of this analysis is to create a linear model of SAM temporal auditory processing by quantifying increases in evoked single-unit firing rates in response to decreased SAM modulation depth.

The linear regression model seeks to estimate a linear relationship between one (simple linear) or more (multilinear) predictor and measured variables. In this model, both the measured result and predictors are metric, as opposed to categorical, variables which map to a continuum of possible values. The simple linear regression model takes the form of:

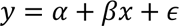

where *y* is the measured (predicted) group, *x* is the predictor, β is the “slope” parameter dictating the relative increase or decrease in *y* per unit change in *x*, α is the intercept term which, in models of firing rate represents non-evoked, spontaneous firing rates, and ε is an error term which quantifies the difference between the expected value of *y* at a given *x* given a linear model versus the observed value of *y* at*x*. It should be noted that εε is not present in all regression models, but the authors suggest inclusion to quantify deviations from linear fit.

Linear regression thus forms a model in which SAM depth predicts evoked firing rates in which the model parameters are estimated and used to draw conclusions about the relative dependency of *y* on *x*. To begin, an observation of the relative distribution of the measured data, in this case firing rates elicited from IC, will allow for robust inference model design. Data distributions are most often visualized through construction of data histograms. Probability density functions (pdf) can then be estimated through kernel density estimation (KDE), a process in which data points are convolved with gaussian kernels to create a smooth estimation of underlying continuous data pdfs(Rosenblatt, 1956; Parzen, 1962). Inspection of the distribution of firing rates (Fig 2B) suggests that a log transform would allow for the data to be normally distributed, making model computations easier through use of canonical normal distributions.

### Performing Bayesian Inference on the Linear Regression Model

Turning back to the example of IC single unit firing rates in response to SAM depth stimuli, the first step in inference is to place a prior distribution on the data. Previous studies and data can be used to inform the prior, but for this example we chose to demonstrate regression with moderately informative priors on α, β, and ε so as to let observed data drive posterior inference. Given that the observed data is roughly normal, a good first pass is to place a normal distribution on the prior with mean equal to the mean of the observed data and a variance that is wide enough to capture all observed data. After inference is made, sensitivity analyses can be performed to assess the relative importance of the prior parameter values on posterior estimates. Larger prior variances allow for small, but non-zero probabilities on extreme values. This tends to be a more robust approach than setting a value of 0 on extreme events, as observed data with strong evidence for an extreme value can be adequately represented in the posterior. After observation of the underlying distribution of the observed data and decision on a prior distribution, a linear regression inference model can be easily described in code as follows:

#### Code Example 1: PyMC initialization of a simple linear regression model

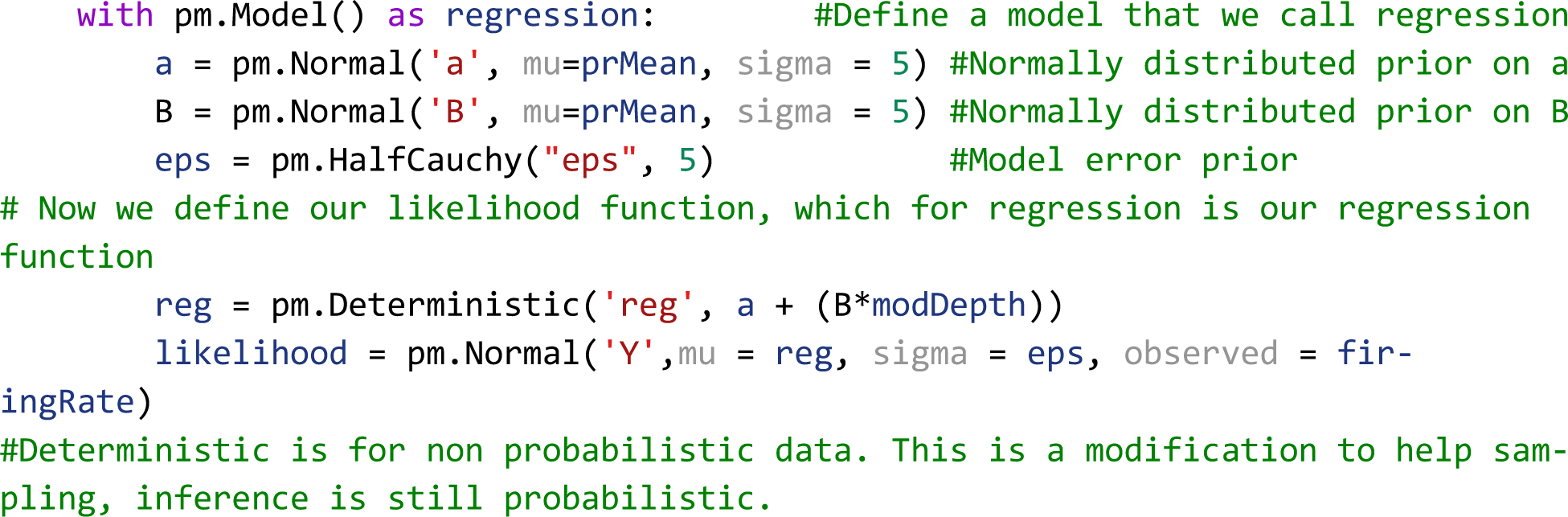

The likelihood variable then translates our model to one of Bayesian inference by casting the model as a probability distribution, in this case

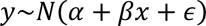

Importantly, Bayesian inference does not require any extra preprocessing of the data outside of what is dictated by the experimental design. In this case, we are interested in estimating assessing the interplay of varying modulation depth on maximum firing rates. Our raw data then is estimated mean firing rates calculated from PSTHs, which are incorporated into the regression model in the “observed” variable in the likelihood function. To generate the posterior, all that needs to be done is to initialize and run MCMC as follows:

#### Code Example 2: Running the MCMC sampler

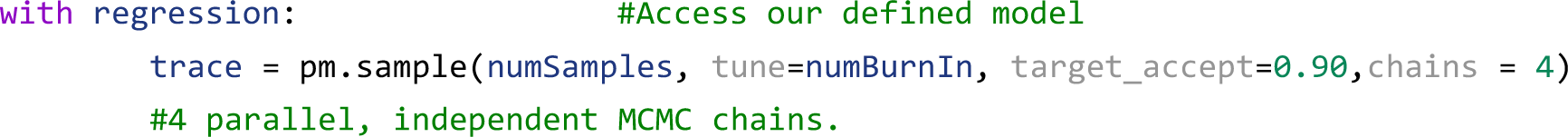

This routine calculates the regression model, generating a trace variable containing the posterior distributions of all model parameters after sampling numSamples with numBurnIn samples to initialize chains. We also ran 4 chains in parallel with a target_accept probability of 90%. Acceptance probability is somewhat based on the statistics of observed data and model, with more difficult posteriors benefiting from higher accept probability values(Gilks et al., 1996). Improper acceptance probabilities can give rise to insufficient number of draws and malformation of posterior distributions. PyMC provides a helpful readout for when posterior draws are malignant and indicative of higher acceptance probabilities. In summary, in a few lines of code the researcher has observed distributions of the data and explicitly defined a model of the data generator and likely now has a better intuition of the data and how it is distributed. All that’s left to observe the posteriors with HDIs to infer significance from the model.

Plotting the 95% HDI estimation of the regression line (Fig 3A) of modulation depth vs natural log-transformed firing rates suggest a small but significant increase in firing rates with increases in modulation depth as evidenced by a 95% HDI credible region (0.015-0.022) that does not include zero from the β posterior distribution (Fig. 3B). A MAP value of 0.018 represents the most probable slope value from the regression. Posterior distributions of model parameters (Fig. 3B) also show that there is an estimated basal firing rate above 0 (α MAP = 3.1) with model error terms considered small for being significantly smaller than intercept term (∈ MAP = 0.74). The spread of the 95% HDI on inferred parameters is used as a measure of uncertainty of the parameter, with narrow HDIs representing more certainty in MAP estimated parameter. In our model, the αα parameter has a spread between 3.02 to 3.13, with a difference of 0.11 containing 95% of its posterior distribution, suggesting strong certainty in the MAP estimate of 3.1. Similar narrow spread is see in the β parameter, with a difference of 0.007 containing 95% of the posterior. The model error term shows that observed data deviation from the model is constrained between 0.71 and 0.76 suggesting relative certainty in the magnitude of deviation of the data from the model. In the context of regression, these posterior results can be interpreted as a mean firing rate increase by a factor of 1.02 (2%, *e*^0.018^) per percentage change in SAM depth that follows a most probable model of

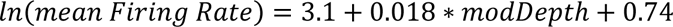

**Figure 3:**
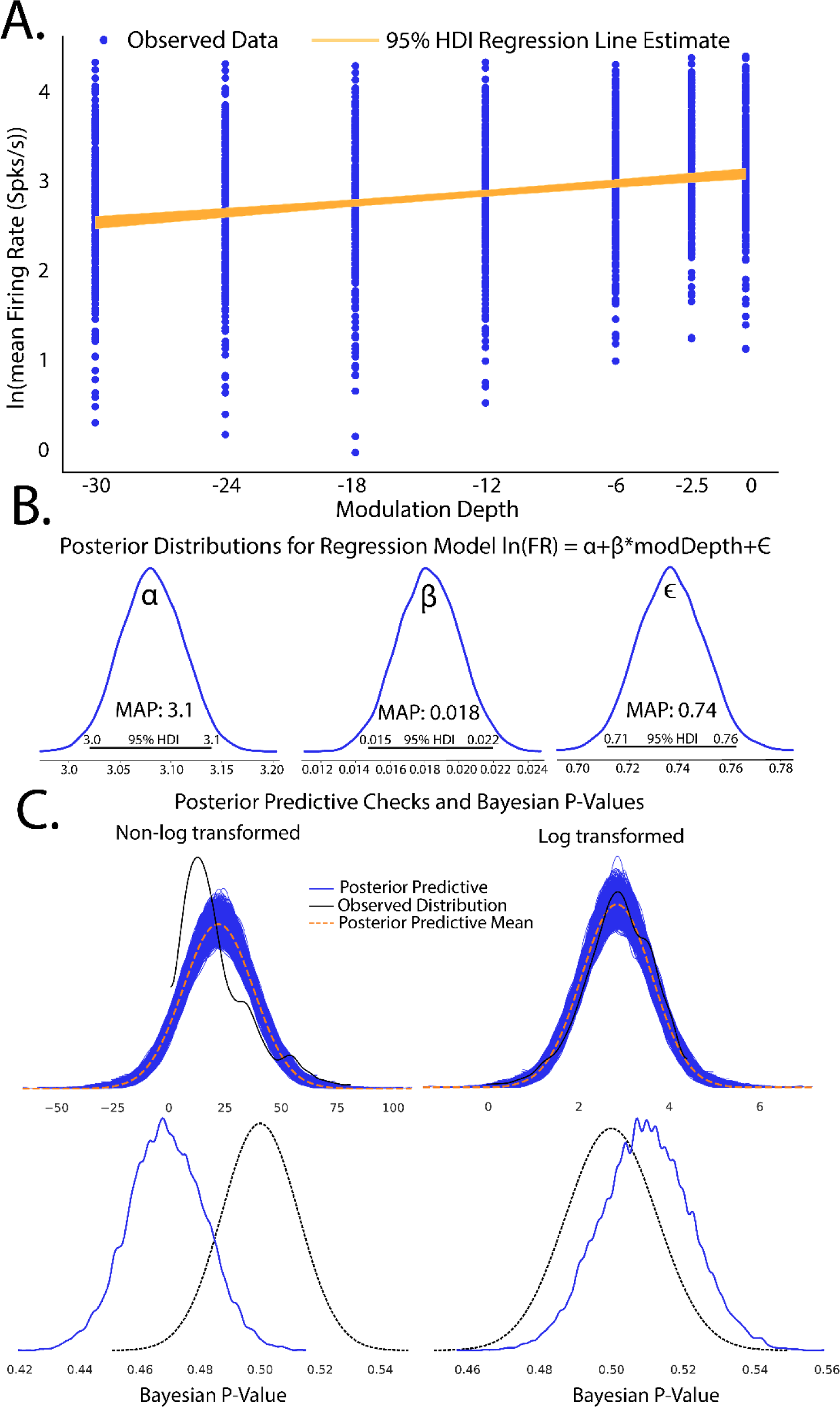
Completed Bayesian inference quantifying linear relationships in evoked firing rate from increases in modulation depth. A. Scatterplot of observed firing rates vs SAM depth stimuli with fitted regression line estimates superimposed. 95% HDI estimates of regression slopes are shown in orange, with the spread of lines encoding the 95^th^ percentile of most likely slope values. B. Estimates of Bayesian linear regression parameters. Intercept term αα was significantly above 0 (MAP = 3.1, 95%HDI does not overlap 0) which indicates basal firing rates above 0. Regression slope was small but significantly above 0 (MAP = 0.018, 95% HDI does not overlap 0) suggesting an increase in evoked firing rates with increased modulation depth. Error term εε was significantly above 0 (MAP = 0.74, 95% HDI does not overlap 0) suggesting some model deviation from observed data. However, error terms were considered small as εε MAP < αα basal firing rate MAPs. C. Posterior predictive checks of linear (left) and log linear (right) regression models show that log transformed firing rate models produce posterior predictions most inline with observed data. Disparity of empirical posterior predictive distributions from observed data as quantified through Bayesian P-values also suggest log transformed firing rates creates a superior model fit.

The ∝ term in the context of this study represents the natural log firing rate for 0% modulation, corresponding to a pure tone stimulus. Statistical conclusions should not end after making inferences on model parameters however. Critical to the validity of statistical inference is the quality of the model fit to observed data. This goodness of fit in Bayesian approaches can be analyzed by posterior predictive checks, in which sample draws are made from the posterior distribution, simulating observations of data generated from the experiment from which the statistical model was fit, and comparing sampled to observed data to assess deviation of model predictions from observed data distributions. In PyMC, posterior predictive checks can be easily performed using the following code:

#### Code Example 3: Performing posterior predictive checks

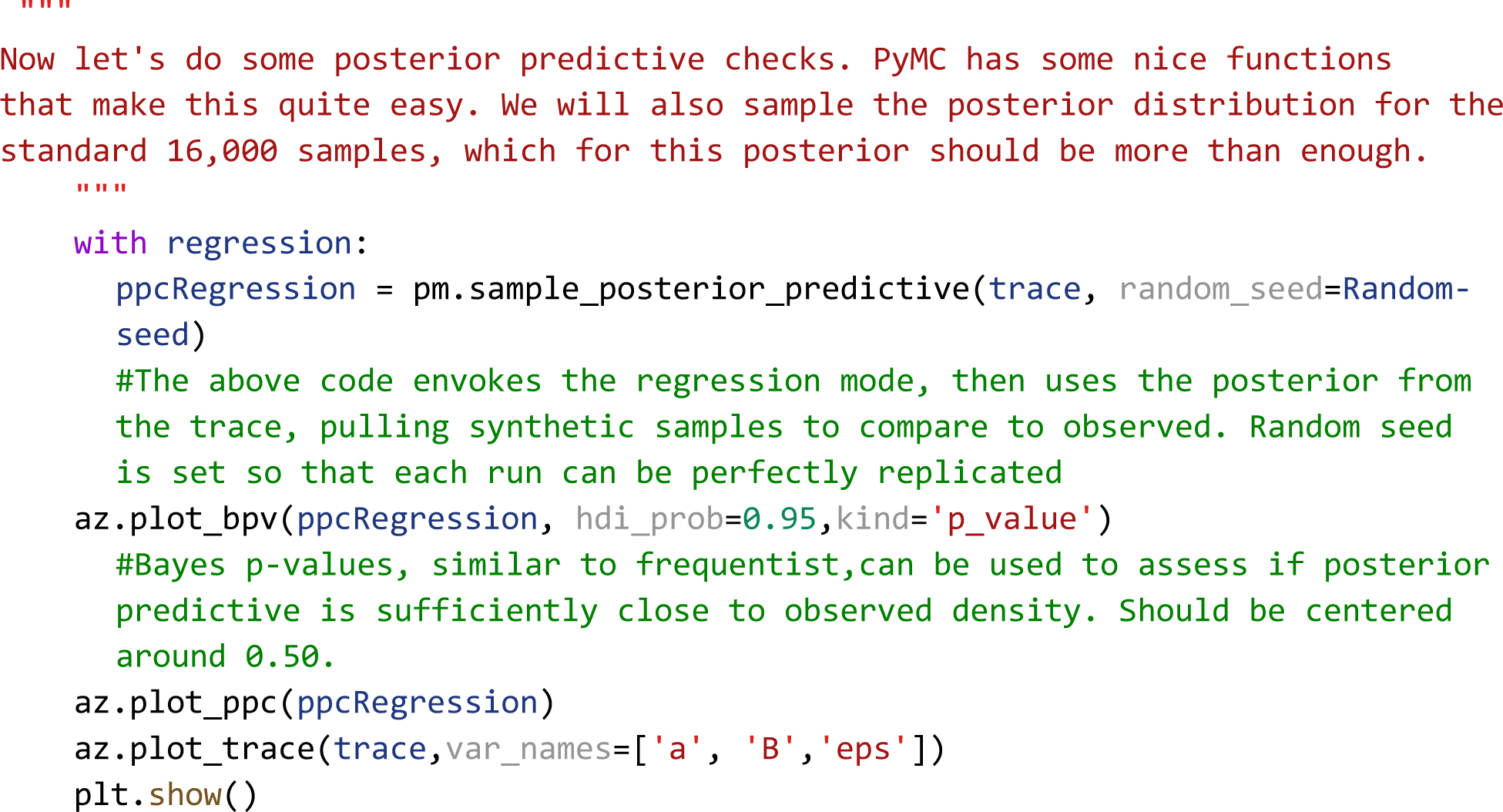

To illustrate how posterior predictive checks can be used, a competing model was made which performs Bayesian linear regression to the same data and priors except without log transformation of the data. In each case, random draws were made from each log transformed and non-log transformed posteriors to create empirical data distributions. Comparison of empirical distributions qualitatively show that log-transformed models present a better fit to observed data than non-log transformed models. The relative disparity between posterior predictive model fits and observed data can be quantified by use of Bayesian p-values(Fig 3C), a distance measure between two distributions (for details of Bayesian p-values, see Kruschke, 2014). The closer the Bayesian p-value is to 0.5, the better data sampled from the posterior overlaps with the distribution of observed data. Plotting the resulting distributions and the Bayesian p-values indeed show the log-transformed model fits better to observed data than the non-transformed model. Similar analyses can be performed around model free parameters, such as prior variables, to form a sensitivity analysis of our of prior distributions on resulting posterior inferences.

A secondary and quick check of posterior sampling can be performed by qualitative evaluation of the MCMC sampling chains, often called traces. Traces represent the long term run of a Markov chain which represent the distribution of interest. As such, good traces show evidence of effective sampling and convergence to target probability distributions. PyMC offers easy ways to visualize posterior MCMC traces using the *plot_trace* function. Figure 4 shows traces obtained from our Bayesian regression example. Kernel density estimates of traces corresponding to the posterior distributions of regression parameters show good convergence of MCMC traces to a target distribution (Fig 4A). As MCMC chains are time series samples which form a distribution, evaluation of traces through sampling time can also be used as a diagnostic of sampling convergence. Traces should have a “fuzzy caterpillar” like appearance (Fig 4B) without any stark jump discontinuities from sample to sample. Quantitative trace evaluations are also available, with the Gelman-Rubin statistic (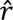) being the most prominent. The Gelman-Rubin statistic measures the variance between MCMC chains to the within chain variance, effectively measuring chain stationarity and convergence(Gelman and Rubin, 1992). Heuristically, 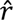 < 1.05 is considered good convergence of MCMC chains. This value can be calculated *post hoc* after sampling and PyMC will automatically flag if 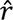 ≥ 1.05 is detected.

**Figure 4:**
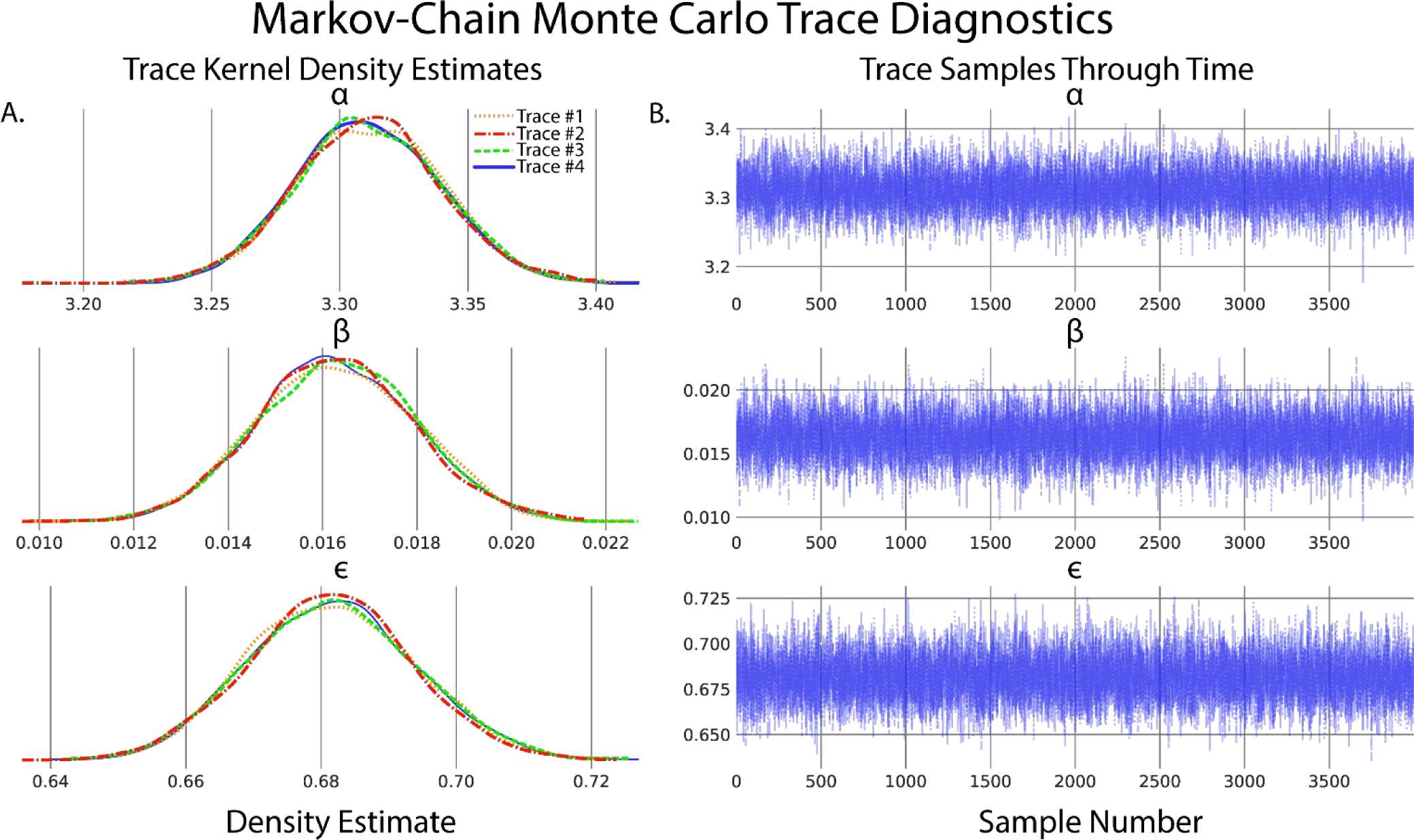
Evaluation of Markov-chain Monte Carlo (MCMC) chains can help diagnose ill-fitting distributions. A. Kernel density estimates of the marginal posteriors corresponding to each of the regression parameters of each MCMC trace. Qualitatively, chain distributions should appear similar to each other, suggesting good convergence to target distributions. B. Time series plot of trace value vs sample number of marginal posteriors corresponding to each regression parameter. Qualitatively good traces should have a “fuzzy caterpillar” like shape, evident in all parameters of this model, indicative of good integration over the joint posterior distribution and effective sampling of the posterior.

### Inference Reporting Guidelines

Proper presentation of statistical methods and processes is critical to the interpretation of conclusions drawn from inference. While there are many reporting guidelines for Bayesian inference, we follow the Bayesian Analysis Reporting Guidelines as given by Kruscke(Kruschke, 2021) and provide an example reporting document including posterior predictive checks, Bayesian model comparisons, and sensitivity analysis in our Github code repository.

### T-Tests and Comparison of Groups

Comparison of differences between two or more groups is one of the most fundamental inference paradigms, ubiquitous to all fields of neuroscience. Frequentist implementations of group comparisons, such as the t-test and χ^2^-tests, suffer from similar ailments as frequentist regressions: strict assumptions about distributions of observed data, lack of interpretability of group differences, and the inability to confirm a null hypothesis. Added to these is that of the multiple comparisons problem, with increasing number of comparisons leading to increasing type-I errors. Bayesian implementations of group comparisons provide complete descriptions of group differences in means, variances, and effect sizes, the ability to confirm and not simply reject null hypotheses, and complete posterior distributions, which remove the need for multiple comparisons corrections(Gelman et al., 2009; Kruschke, 2011, 2013). One implementation of Bayesian group comparisons is Bayesian estimation Supersedes the t-Test (BEST)(Kruschke, 2013), consisting of a hierarchical model with data described by t distributions and minimally informative priors on group means and standard deviations.

### Comparison of β-Band Dynamics in a DBS Model Using BEST

To illustrate the use of BEST in group comparisons, a computational model of the dopamine-depleted basal ganglia thalamocortical circuit was utilized (Fig 5A). Simulations consisted of measurements of oscillatory activity in local field potential (LFP) β-band (13-30Hz), elevation of which is a known biomarker for Parkinsonian symptomology across the basal ganglia circuit(Singh, 2018). “No stimulation”, in no stimulation, subthreshold “sham” stimulation, and effective DBS stimulation groups were tested. LFPs were recorded from globus pallidus internus and stimulation in subthalamic nucleus with 2000 repetitions of each group performed. Figure 5B shows example LFP power spectrums for no stimulation, subthreshold stimulation, and DBS conditions respectively, with reductions in β-band activity shown in DBS conditions compared to no stimulation and subthreshold stimulation respectively. Posterior distributions allow for estimates of group means and standard deviations of LFP β power (Fig 5C) with qualitative observations of non-overlapping credible regions of means (DBS: 0.46-0.47, No stimulation: 0.86-0.88) and standard deviations (DBS: 0.036-0.041, No stimulation: 0.068-0.078) suggesting that DBS produces strong reductions in β power. Unlike frequentist methods, direct measurements of differences in group means are easily quantified from subtraction of group posterior distributions. We reiterate that unlike frequentist methods, this inference is formed directly from observed data, and not conditioned on hypothetical population distributions. Differences in group means show strong reductions in β-band power in DBS conditions (MAP estimate μ_*DBS*_ − μ_*nostim*_ = −0.4) with MAP estimates of effect size showing strong, discernible evidence of DBS effect in reducing β oscillatory activity (MAP = -6.9). Differences in group means and effect size are considered significant because 95% HDIs do not contain 0.

**Figure 5:**
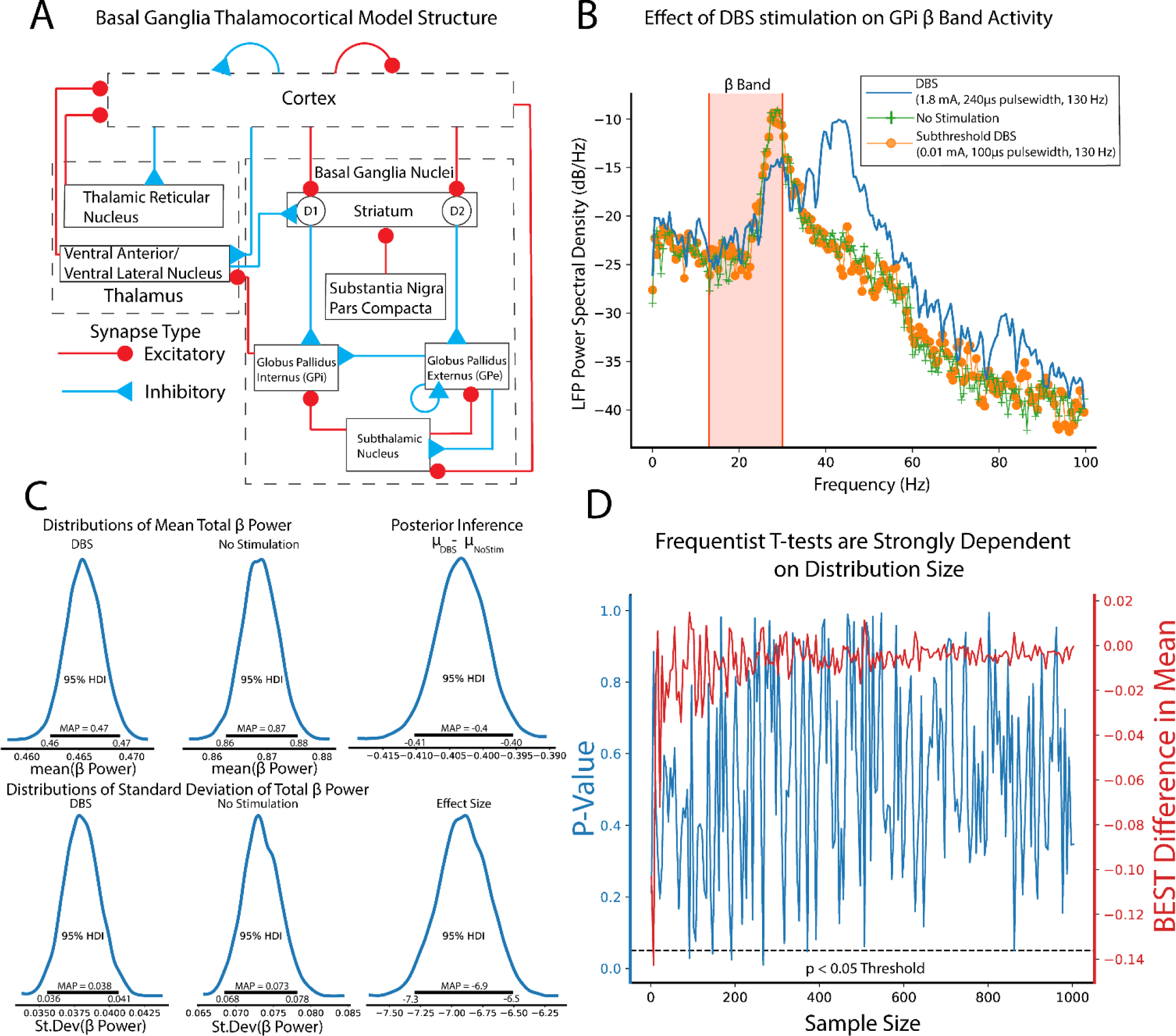
Bayesian implementations of t-tests provide descriptive quantification of differences in group means and standard deviations and effect sizes drawn directly from observed data. A. Schematic of the basal-ganglia thalamocortical computational model. Modeled DBS stimulation was performed in subthalamic nucleus with local field potential (LFP) recordings made from globus pallidus internus. All simulations were performed assuming dopamine-depleted conditions. B. Example LFP power spectral densities from no stimulation condition (green), subthreshold DBS stimulation (0.01 mA, 100 μs pulse widths, 130 Hz, orange), and effective DBS stimulation (1.8 mA, 240 μs pulse widths, 130 Hz, blue). C. Total β-band power in no stimulation and DBS stimulation was calculated (n=2000 repitions). Group comparison by Bayesian estimation supersedes the T-test (BEST) shows significant reduction in mean β -band power as evidenced by group mean difference 95% HDIs not including 0 (MAP μ_DBS_ – μ*_nostin_* = −0.4). Estimated effect size distributions further give evidence of a strong DBS effect of lowering β -band power in dopamine-depleted conditions (MAP *_effectsize_* = 6.9, 95% HDI does not contain 0). D. Frequentist t-tests show strong dependance on sample size. Comparisons of β-band power were made between no stimulation and subthreshold stimulation groups. Subthreshold stimulation will have no effect on β-band power with only differences formed by model noise simulating intrinsic neural noise. Increasing numbers of shuffled samples were included in t-test and BEST tests. BEST differences in group means show rapid convergence to null differences. P-values from frequentist t-tests however show drastic fluctuations some of which cross *p* < 0.05 significance threshold even at large sample sizes.

Estimation of group differences further shows the explanatory power of Bayesian inference in its ability to provide direct, probabilistic estimates of group summary statistics, group differences, and effect sizes. Bayesian approaches with appropriate priors are also more robust to sample size, particularly lower sample sizes that are typical of many neuroscience experiments(Smid et al., 2020). To illustrate this, we utilized frequentist t-test and BEST inference, comparing no stimulation and subthreshold stimulation groups. BEST and t-test estimates on these groups should confirm a null hypothesis/ fail to reject the null hypothesis respectively given no effect of subthreshold stimulation on LFP β -band activity. In this study, random study stopping points were simulated by performing BEST and t-test estimates on increasing sample sizes (2-1000, steps of 5). Data points were pulled from random permutation of no stimulation and subthreshold stimulation groups. BEST estimation showed rapid convergence to estimated group differences of 0 at low sample sizes, while frequentist p-values showed strong fluctuations regardless of sample size (Fig 5D). P-value oscillations showed several instances of falling below *p* < 0.05 decision criteria, even at sample sizes larger than the majority of neuroscientific experiments. While p-value dynamics are contextual, based on data type and underlying data distributions, this data shows the potential of NHST to draw drastically different experimental conclusions from sampled data distributions with very similar sample sizes.

### Multilinear Regressions, Repeated Measures, and Hierarchical Models

In many experiments, inference across multiple possible data generating parameters must be analyzed and accounted for. These models, called multilinear regressions, are extensions of standard linear regression as follows:

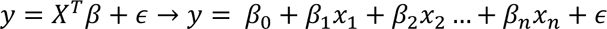

where n is the total number predictors.

To illustrate the use of multilinear regressions, consider the case of thalamocortical infrared neural stimulation (INS)(Fig 6A). Auditory thalamic neurons in the medial geniculate body were excited by pulse trains of optical stimuli varying in pulse energy and time between pulses. The resulting auditory cortex single unit responses are recorded using a planar, Utah style array in layer 3/4. An important and understudied aspect of INS is the effect of laser energy and interstimulus interval changes on evoked firing rate responses; a so-called dose-response curve. We begin by specifying predicted and predictor values. Dose-response relationships were measured by predicting maximum firing rates in response to applied INS energy (E) and inter-pulse intervals (ISI). As we suspect an interaction between E and ISI, an interaction term of E*ISI was incorporated. Therefore, the model was defined as:

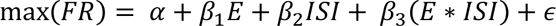

**Figure 6:**
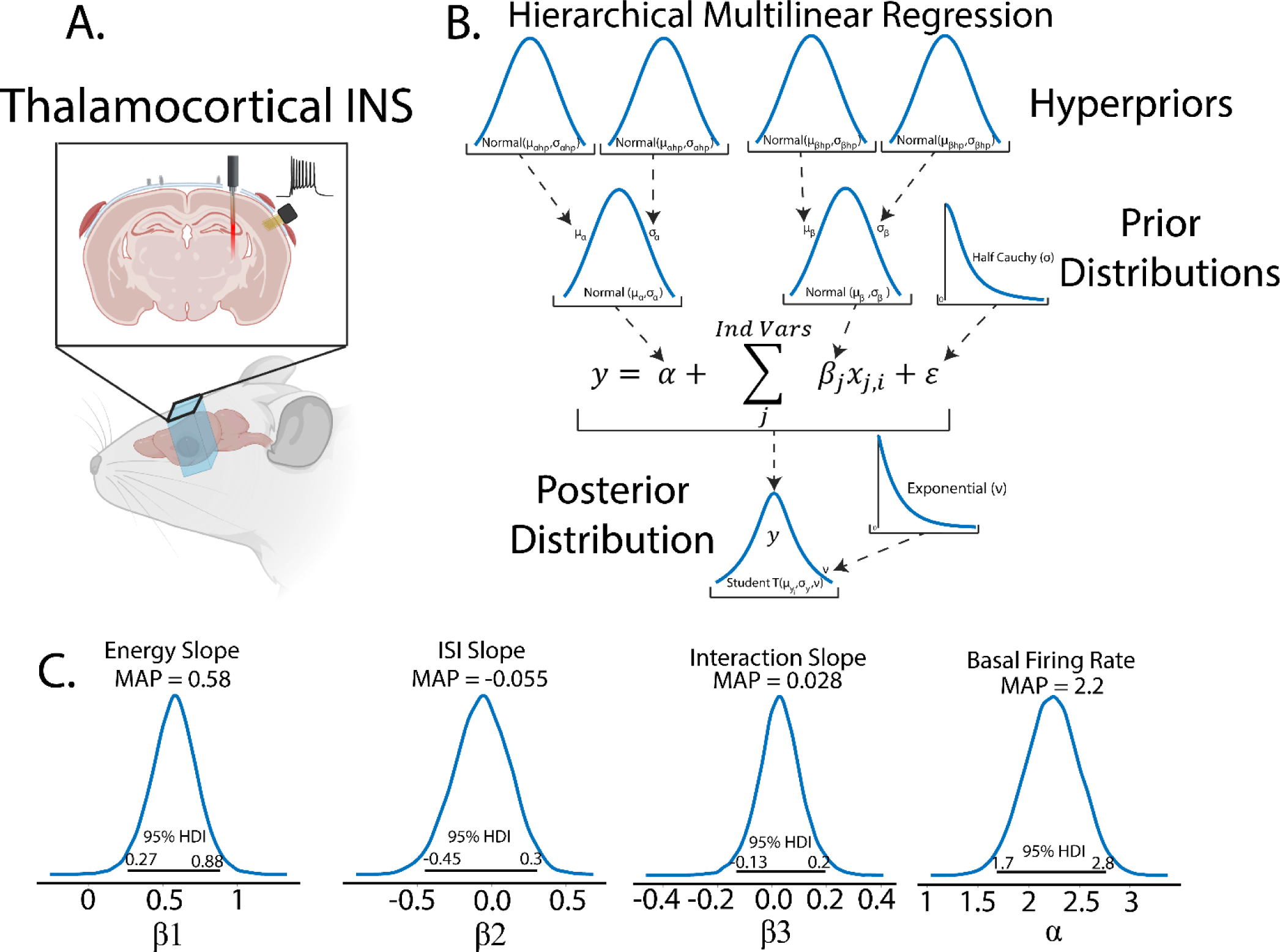
Example of Bayesian multilinear regression incorporating a hierarchical structure. A. In this experiment, rodents were implanted with fiber optic arrays into auditory thalamus and planar recording arrays into auditory cortex. Single unit responses were recorded from INS stimuli with applied energy and interstimulus intervals varied to derive dose-response curves. Figure was drawn using BioRender under publication license (www.biorender.com). B. Hierarchical schematic of Bayesian multilinear regression. Hierarchical structures are advantageous in accounting for within and between subject variability or for repeated measures designs. C. Resulting parameter distributions from dose-response models. Energy was a significant contributor to maximum firing rate, with increasing laser energy resulting in increased maximum firing rate, as determined by 95% HDI of the laser energy term β_1_excluding 0 (MAP = 0.58). Laser pulse interstimulus interval did not significantly contribute to changes in max firing rate as indicated by ISI parameter β_2_overlapping 0 in its 95% HDI with a MAP value near 0 (MAP = 0.028). The relatively wide spread about zero does suggest that there may be a subset of ISIs which contribute more strongly to firing rates and warrants further study. Laser energy-ISI interactions also did not significantly contribute to max firing rate as evidenced by interaction parameter β_3_including 0 in its 95% HDI. The intercept term αα, correspondint to basal firing rates, were significantly above 0 (MAP = 2.2, 95% HDI excludes 0).

An important aspect of this study was that rats underwent chronic recordings through the duration of the lifetime of the implant. It almost a certainty that stimulation and recording quality will change over the lifetime of the devices due to neural adaptation to stimulation(Falowski et al., 2011) and glial response and encapsulation of the devices(Van Kuyck et al., 2007; Woolley et al., 2013). This experimental paradigm is thus complicated by potentially meaningful repeated measures within subject variability. Furthermore, slight differences in electrode and optrode placement between rodents could create a heterogeneity in the receptive fields of recorded neurons(Vasquez-Lopez et al., 2017), representing a potentially meaningful between-subject variance.

### Hierarchical Structures Capture Latent Variables

Models in both Bayesian and frequentist paradigms capture these within and between subject variances by adding hierarchical structure to the model. From the Bayesian perspective, hierarchical models are defined by allocating hyperparameters on the prior which encode within and between group variances in the model, with each hyperparameter containing hyperprior distributions. Graphically, this is organized in Fig 6B. Bayesian and frequentist hierarchical models share similar roots, with particular hyperprior distributions in Bayesian paradigms becoming proportional to frequentist random effects models.

While this appears to be a herculean task in data modeling, PyMC allows for declarations of hierarchical models, as shown in Code Snippet 4:

#### Code Example 4: Creating a hierarchical regression model

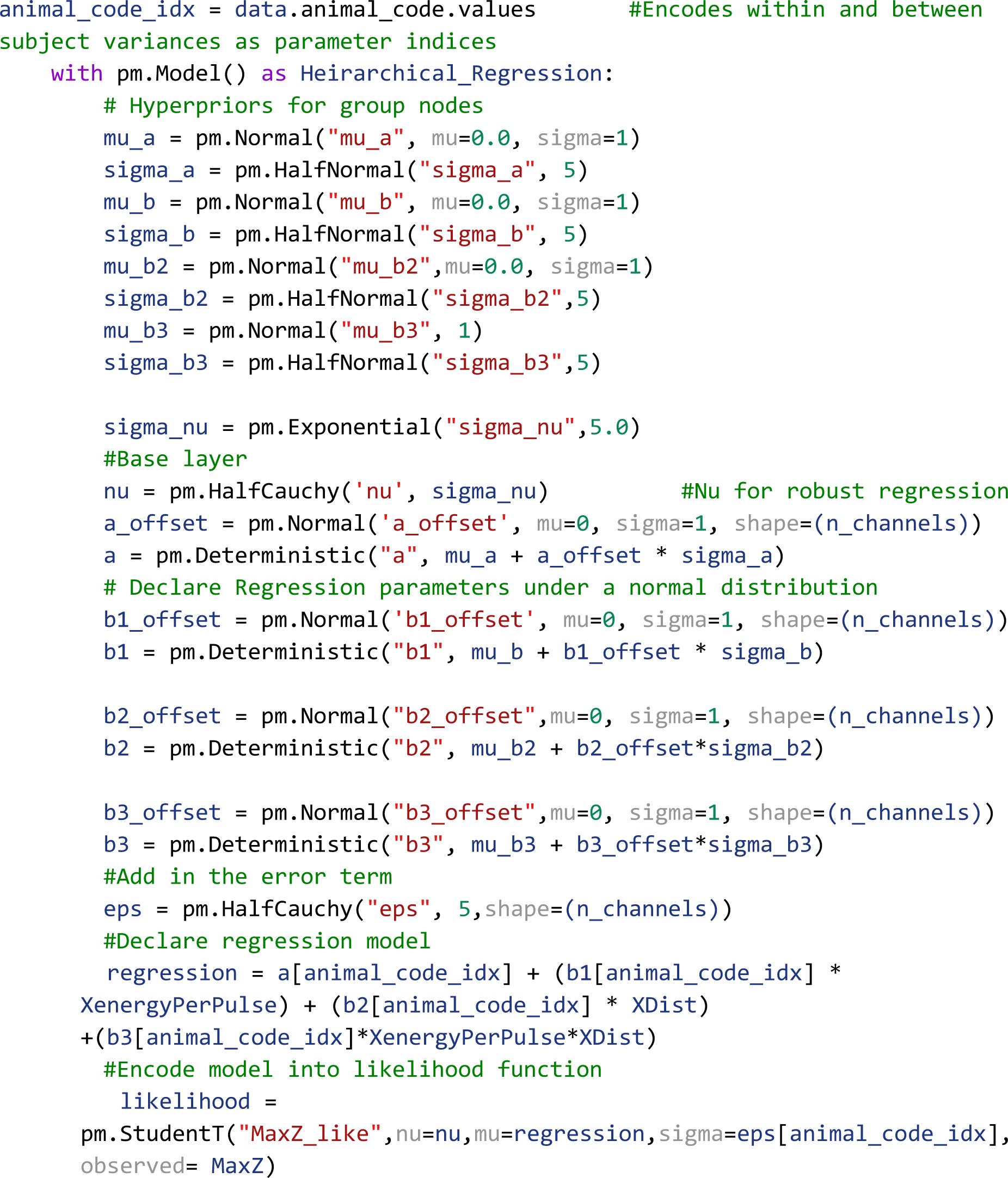

Owing to the scarcity of thalamocortical INS data, we assigned noninformative, wide spread normal distributions on the priors and hyperpriors so as to let the data speak for itself. We also utilized a student-T distribution as the likelihood function to accommodate outliers in a modification known as “robust regression”(Kruschke, 2014). Student-T distributions have tails which are not bounded by the exponential function, meaning that extreme values have less impact or skew on the posterior distribution. Half-Cauchy distributions are placed on the error term and Student-T normality parameter *v*. Half-Cauchy distributions are advantageous in learning scale parameters from the data in hierarchical models (Gelman, 2006; Polson and Scott, 2012).

It is important to validate that our model and data generating functions indeed represent the observed data. Sensitivity analyses and posterior predictive checks thus can be performed to ensure the model chosen is the one that best describes the observed data. Sensitivity analyses were performed by varying prior variance and comparing models which were nominal or natural log transformed with normal and student-T likelihood functions. Model comparisons can be performed in many ways, but a common paradigm is the leave-one-out cross validation (LOO)(Gelman et al., 2014). LOO consists of partitioning data into training and test sets and iteratively fitting the model under test with training data and testing out of sample fits with test data. Models are then ranked using the expected log pointwise predictive density (ELPD) measure:

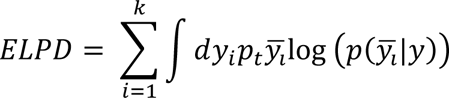

where *p*_*t*_, *y*_*i*_ are unknown distributions representing the true data generating function for estimates of true posterior predictive function 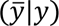 from observed data *y*(Vehtari et al., 2017). In general, larger values of ELPD represent better out of sample fits indicative of a better model conditioned on observed data. We can then use standard errors between the model with the best ELPD (dse) and all competing models to rank all models to observed data. Importantly, these metrics should be understood only in the context of a model relative to other models, and not a global predictor of model validity. Observations of posterior fits to the data using posterior predictive fits and Bayesian p-values should be utilized on the final model to determine model fit. This seemingly complex model comparison can be quickly and easily done in PyMC with the following commands:

#### Code Example 5: Model Comparisons

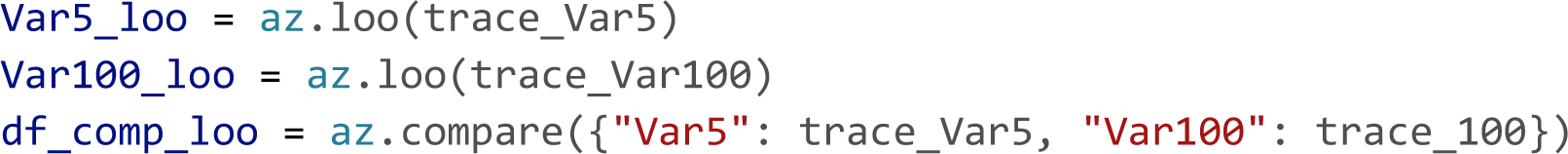

Model comparison results are given in Table **2**. Similar to the simple regression above, the log transformed model provided much better fits to observed data than non-log transformed models. Interestingly and instructively, moderately informative priors (variance 5) outperformed noninformative priors (variance 100), suggesting that constraining prior variance can have predictive power in inference. Posterior predictive checks on the winning model show good fits to observed data with a Bayesian p-value near 0.5.

**Table 2:**
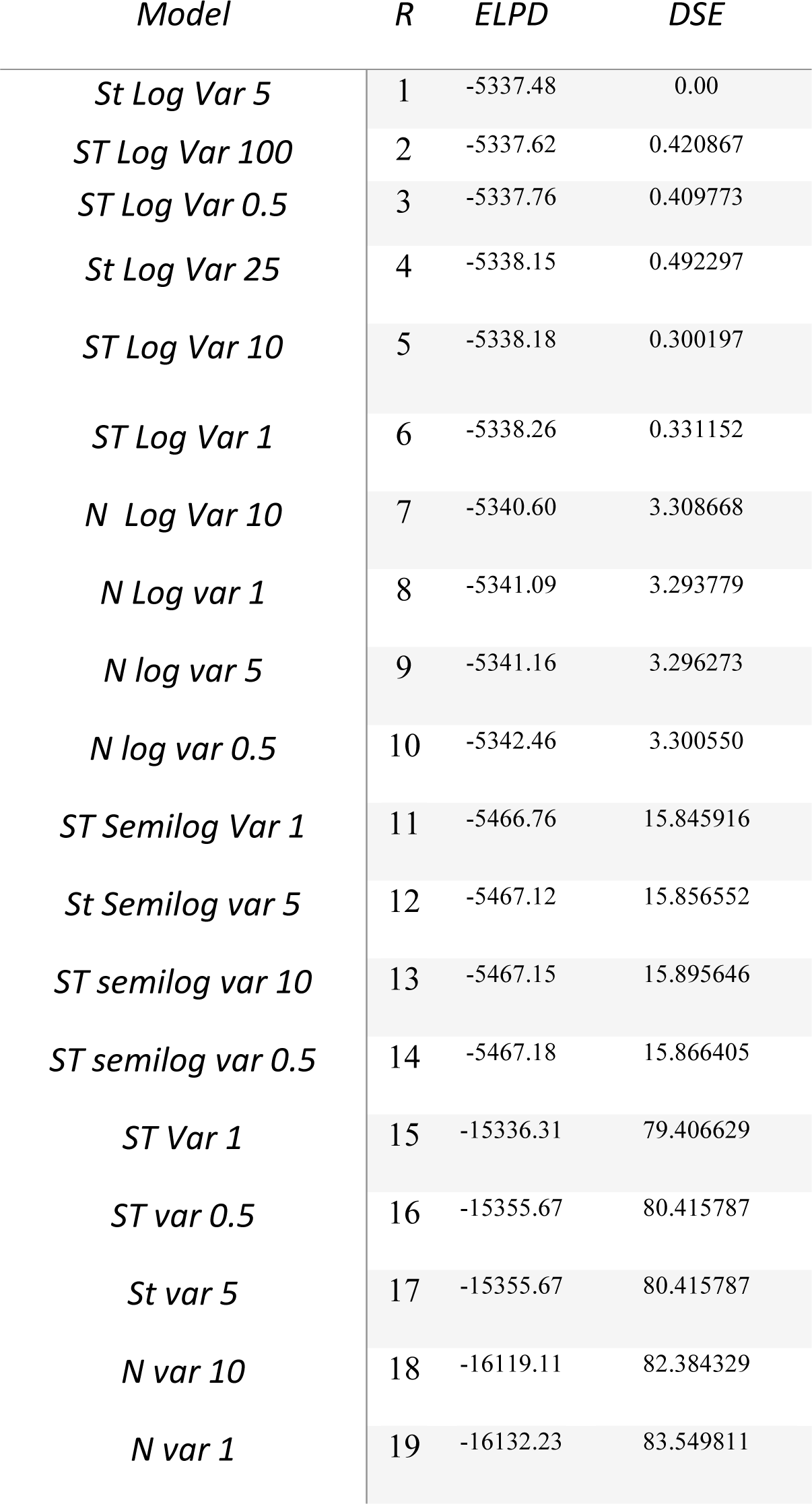
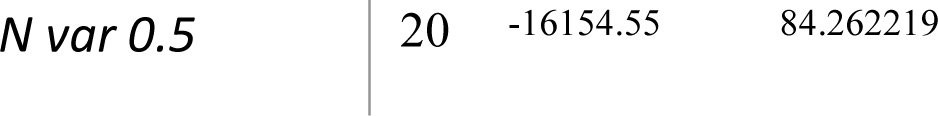
LOO Model comparisons and sensitivity analyses.

We can now perform inference on our multi-regression model. It was found (Fig 6C) that αα was significantly above 0 (MAP = 2.2, 95% HDI does not cross 0) suggesting that basal firing rates of recorded neurons were typically above 0 as expected. It was also seen that maximal firing rates were significantly dependent on applied INS energy (β_1_MAP = 0.58, HDI does not cross 0) with increases in INS energy leading to larger evoked maximal firing rates. The relative spread of the 95% HDI on ββ_1_of 0.27-0.88 suggests a heterogeneity in neuron dose-response characteristics that can be explored more. Somewhat surprisingly, there was no significant effect of ISI on maximum firing rates (β_2_ MAP = - 0.055). The relative spread across 0 of -0.45 to 0.3 suggests that extreme values of ISI might potentially have an effect, with smaller ISIs causing neural integration of singular INS pulses into a singular, large pulse. However, that cannot be determined given the INS parameters used in this study. Also surprisingly, there was no significant effect of Energy-ISI interactions (β_3_ MAP = 0.028), suggesting that INS energy is the primary mediator of evoked firing rates.

### Bayesian ANOVAs

Comparison of differences between groups is another routine statistical procedure used when predictor variables are nominal or categorical in nature or a mixture of metric and categorical predictors. The frequentist treatment of these experimental designs largely uses analysis of variances methods, namely ANOVA for categorical predictors and, more generally, ANCOVAs for categorical predictors with metric covariates. ANOVAs are models that take the form of:

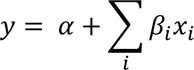

where β_*i*_, *x*_*i*_ are the parameters corresponding to nominal predictor class *i*, α is the offset or bias parameter, and *y* is the metric dependent variable. ANOVA parameters and class values β_*i*_, β_*i*_ are treated differently than the regression case, as β_*i*_ are categorical as opposed to continuous, metric values. As such β categories are recast into “one-hot” encoded vectors 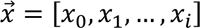 in which only a singular value in an array can have a value of 1 and all other elements are cast to 0, allowing for binary indication of a given class among a group of classes. If an individual value falls into group *j*, for example, 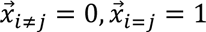. The coefficients β_*i*_ then encodes the change in dependent variable *y* from inclusion of datapoint β in category *i*. Importantly, deflections from baseline are constrained such that ∑_*i*_ β_*i*_ = 0. Both Bayesian and frequentist ANOVA models treat β_*i*_ parameters as group deflections about the baseline level of the dependent variable.

ANCOVA is a modification to the ANOVA model to include a metric covariance term:

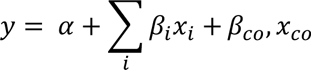

where β_*co*_, β_*co*_ are the parameters corresponding to metric predictors. Metric predictors terms are valuable in accounting for within group variance which is attributable to some other metric measurable variable, such as decreased firing rates in response to an applied stimulus found in a class of aged animals.

Bayesian analogues of ANOVA and ANCOVA can be easily defined in PyMC and are termed BANOVA and BANCOVA (Fig 7A) respectively to distinguish models from their frequentist counterparts. Traditional ANOVAs make two key assumptions; that underlying data is normally distributed and a homogeneity of variance among groups. To account for these assumptions, normal distributions are placed on prior parameter and observed data distributions and a uniform distribution prior is placed on observed data variance σ_*y*_. Importantly, observed data distributions should be assessed to assure distributions are normally distributed. While not strictly an ANOVA-like structure, an advantage of Bayesian approaches is the ability to create models which handle arbitrary distributions. While traditional ANOVAs also assume independent group variances, the relative shared influence between groups can be learned from the data by imposing a hyperprior on group variance σ_β_ (Gelman, 2006). As with any prior distributions, selection of σ_β_ should be informed by prior inspection of the data. A Half-Cauchy distribution is once again chosen as it weakly informative and allows for extreme values if data dictates(Gelman, 2006; Polson and Scott, 2012). Setting σ_β_ to a large constant replicates a traditional ANOVA.

**Figure 7:**
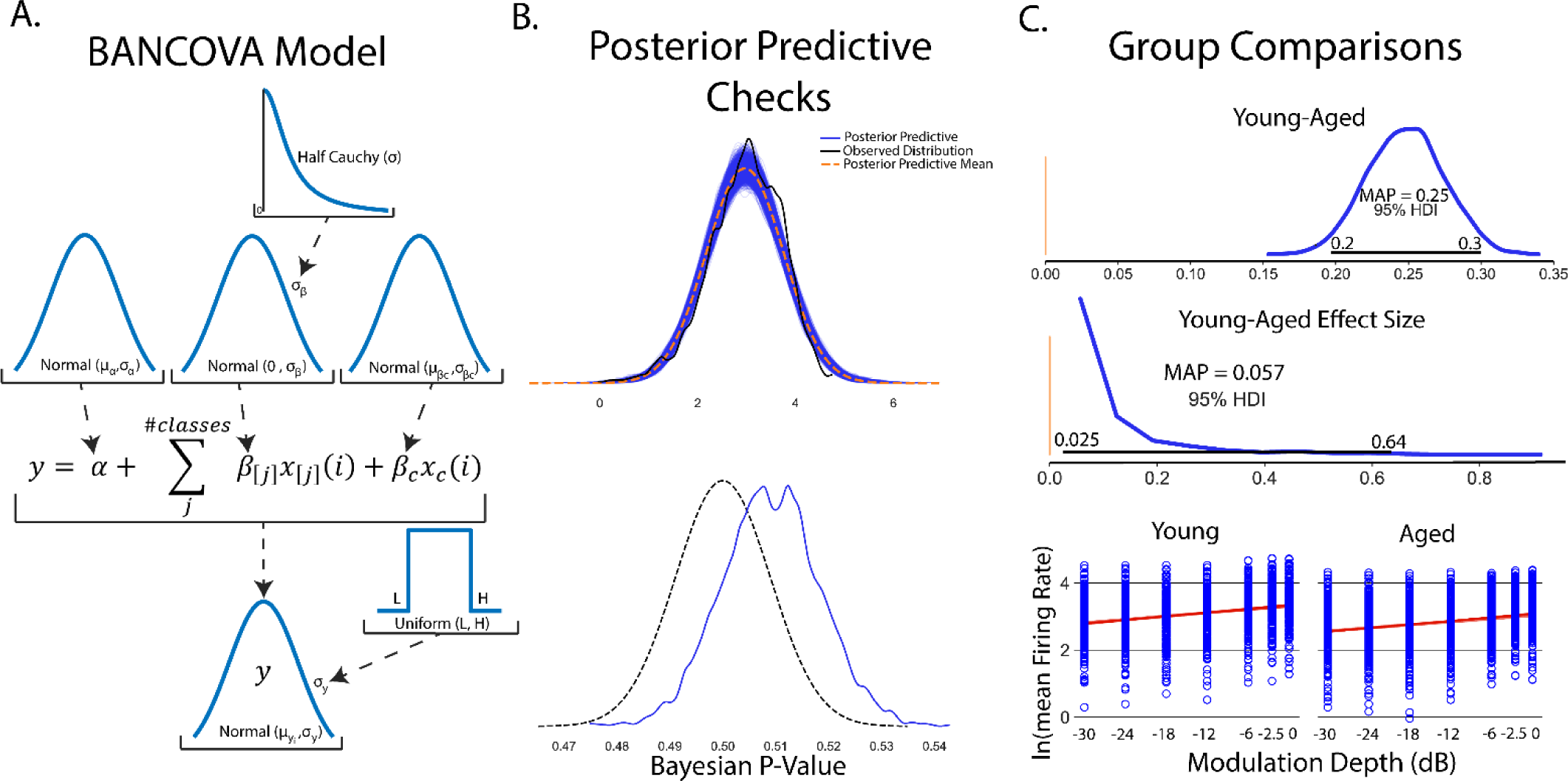
An example of Bayesian inference using ANOVA-like models. A. General schematic of BANOVA/BANCOVA models. Traditional ANOVAs have two key assumptions; normality of group data and homogeneity of variance. Normality of group data is imposed in BANOVA-like models as normal distributions around group parameters with homogeneity of variance encoded as a uniform distribution around posterior variance term σ_*y*_. Traditional ANOVAs assume a fixed variance on group parameter values σ_β_, imposing the constraint that each group is estimated independently from each other group. A uniquely Bayesian approach is to instead learn σ_β_ values from the data itself by placing a distribution on σ_β_. B. Posterior predictive checks suggest posterior distributions show good fit in mean and variance to observed data. C. Once posterior distributions are calculated, group comparisons can be easily done by subtracting young and aged posteriors to yield a contrast distribution. It is found that firing rates across all modulation depths are significantly higher in aged vs young rodents (contrast MAP = 0.25, 95% HDI does not overlap 0). Another unique feature of Bayesian approaches is the ability to assess distributions on effect size. In this BANCOVA, while group differences are significant, their relative effective size is small but significant (effect size MAP = 0.057, 95% HDI does not cross 0) suggesting marginal impact of age on firing rates elicited from SAM stimuli. Finally, metric covariates of firing rate in response to varying SAM depth in young and aged groups can be plotted as regressions superimposed on raw data.

As a guiding example, consider a similar experiment to that done in simple linear regression. In this experiment, we aim to understand age-related changes in IC auditory processing of sinusoidal amplitude modulated sounds. This experiment consisted of two groups of young (animals < 6 months in age) and aged (animals > 22 months in age). SAM stimuli at increasing modulation depths were played to the animals with evoked single unit responses recorded from IC. As seen in the previous simple linear regression experiment (Fig 3), there is a significant increase in evoked firing rate with increased modulation depth in young animals. As such, it should be included in comparison between the two groups. Taken together, this suggests BANCOVA will serve as an appropriate model. BANCOVAs are inherently hierarchical(Gelman, 2005; Kruschke, 2014) (Fig 7A) to allow for between subject variances to be represented in the prior if these variances mutually inform one another. Setting this hyperprior to a constant creates a model analogous to a frequentist ANCOVA(Kruschke, 2014). The formation of the BANCOVA is again relatively straightforward:

#### Code Example 6: Creating a Bayesian ANCOVA

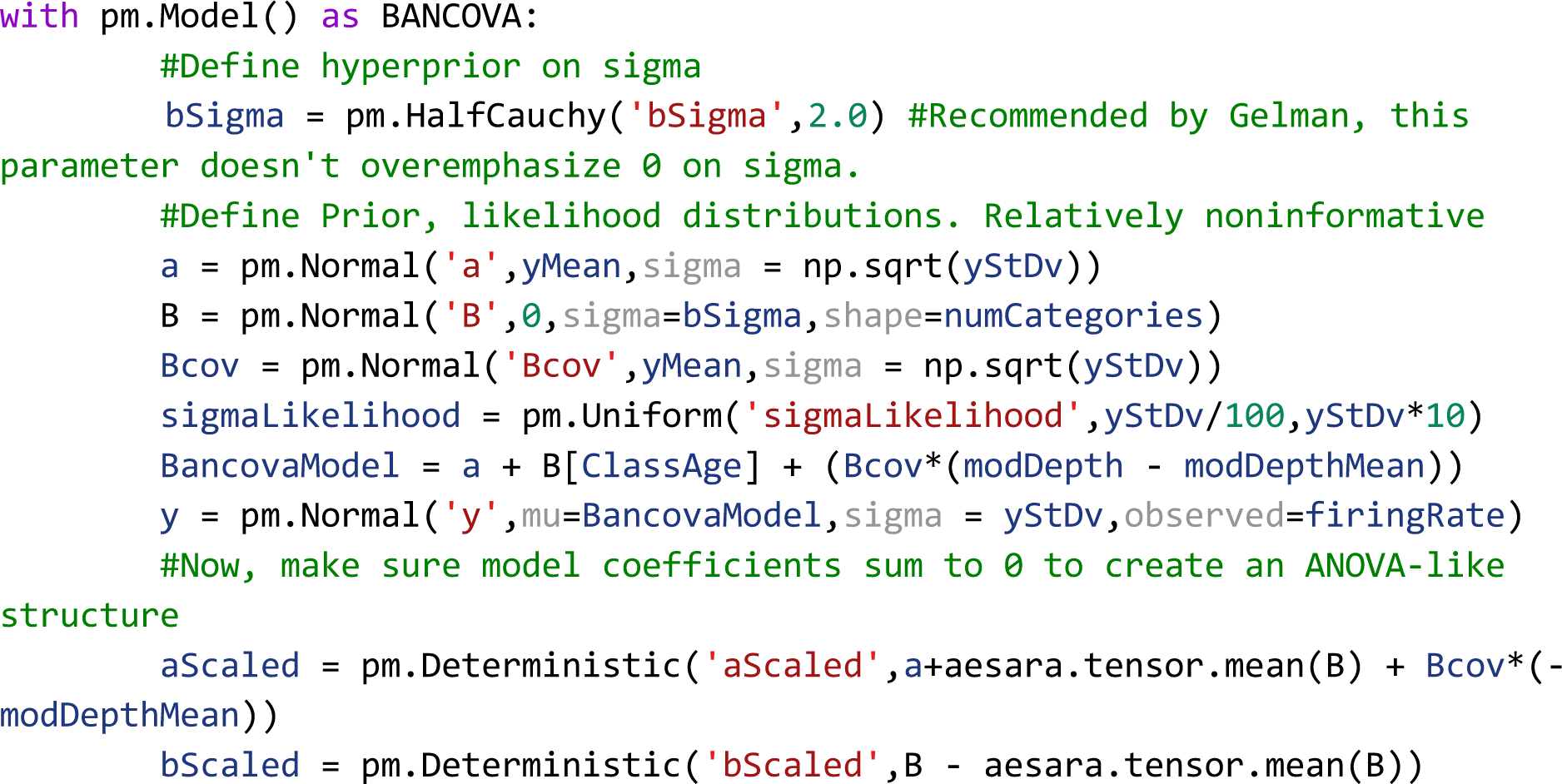

with inference made in the exact same way as the previous models.

After model sampling, posterior sampling checks were performed to ensure posterior distributions adhere well to observed data. Posterior predictive distributions show good qualitative fit to observed firing rate data with Bayesian p-values centered around 0.51, suggesting good model fits to observed data (Fig 7B). Comparisons between groups is simple once posterior distributions are obtained. Similar to Bayesian group comparisons (BEST, Fig 5), all that needs to be done is to measure differences between aged and young group mean parameter posteriors (Fig 7C), encoding influence of young and age groups on firing rates. Aged and young contrasts show significantly elevated firing rates in young rats across all SAM stimuli (Young-aged difference MAP = 0.25, 95% HDI excludes 0). Another advantage of Bayesian inference is the ability to observe the distribution, and thus the most likely value and spread of effect size. In this analysis, the effect of age in SAM stimulus processing is significant but small (effect size MAP =0.058, 95% HDI excludes 0) but with a wide spread of effect (95% HDI between 0.025-0.64) suggesting variable temporal acuity between rodent subjects. Finally, firing rates vs SAM amplitude depth for each class are plotted with *y* = α + β*_young_*_/*age*_*x*_*young/age*_ + β*_cov_x_cov_* superimposed.

### Multiple Comparisons in Bayesian Inference

In traditional frequentist analyses, corrections for multiple comparisons are necessary in order to ensure that maximum Type I errors (false positives) are constrained to a maximum of 5% (α = 0.05). With Bayesian inference, a posterior distribution across all parameters is obtained which remains unchanged no matter how many comparisons are made(Kruschke, 2014). Furthermore, frequentist type I errors are classically defined in the context of rejection of a null hypothesis. Bayesian inference is not strictly concerned with rejection of a null hypothesis, instead weighing competing hypotheses given observed data. Bayesian models are not immune to making false conclusions about data. These errors, called type M for errors in magnitude and type S for errors in sign occur when outliers in data exert too much influence on inference. These errors can be controlled by proper choice of priors or by building hierarchical models (Fig 6A, Fig 7A) which can account for outliers by pulling parameters towards group means when evidence is small and allowing parameters with good evidence to remain in a phenomenon called partial pooling implicit to hierarchical structures(Gelman et al., 2009).

## Discussion

Bayesian inference approaches present a powerful statistical tool which encourages deep and meaningful exploration of data and allows for presentation of data in intuitive and transparent ways. In this tutorial, we demonstrate the ease by which Bayesian inference can be performed across a wide variety of experimental designs and provide source code which can be modified to accommodate neuroscientific experiments using all free and open source tools. We intentionally used the base PyMC toolchain in order to explicitly show Bayesian model creation. However, there are PyMC plugin tools such as Bambi (Capretto et al., 2022) which can facilitate creation of Bayesian models in single lines of code. An example of Bambi-enabled model creation is provided in our Bayesian inference toolbox.

### Applications of Bayesian Inference

In this tutorial, our inference examples largely focused on data commonly found in electrophysiology and computational neuroscience studies. However, Bayesian inference is agnostic to the form and type of data used in inference. The described statistical models are easily adapted to electroencephalography, neuroanatomical measures, behavioral measures, and calcium events, among others. Bayesian inference is of particular interest to neuroscience experiments involving very large datasets, such as spatial transcriptomics (Gregory et al., 2022; Song et al., 2022).

We also focused primarily on canonical statistical model structures of t-test group comparisons, simple regression, ANOVAs, and mixed-effect models. Bayesian models can further be defined for other statistical models such as logistic, Poisson, and ridge regressions, mixture of Gaussians, and time-series analysis.

### Tempering Expectations of Bayesian Inference

Despite the enthusiasm of some Bayesian advocates, Bayesian inference is not a panacea. It is subject to similar problems as frequentist NHST, in that models can be used which do not adequately fit underlying data statistics or priors can be chosen which dominate model performance and deemphasize observed data. However, Bayesian approaches support and encourage model transparency, requiring researchers to declare model priors and posteriors while encouraging continued discussion of inference on data as opposed to stopping if a p-value is below an arbitrary threshold. A second caveat is that running MCMCs can be slower than frequentist approaches, with run times sometimes in minutes as opposed to seconds. However, time increases are not astronomical and can be further reduced to levels similar to frequentist approaches by using GPU computing or using programs such as JASP(Love et al., 2019) which utilize a C backend to speed up computation.

### The Controversy of the Prior

The prior is arguably the most contentious aspect of Bayesian inference, with arguments that the prior unduly influences decisions on data. It is absolutely possible to have priors that distort posterior distributions into poor inference. Similar arguments can be levied at Frequentist approaches which perform similar distortions on decision metrics, such as applying ANOVA tests when underlying data is not normal. Often times, these mistakes are not done out of malevolence, but due to the modern framework of how statistics is performed. We argue that having to consider what prior to use, and thus what one’s assumptions are, what distributions are physiologically relevant, and the distributions of observed data will help to prevent errors in statistical modeling while creating greater transparency in how conclusions on data are drawn.

### Decisions with Bayes Factors

Some studies which utilize Bayesian inference use a decision metric called a Bayes’ factor, which is a measurement of the ratio of marginal likelihoods of two competing models providing log likelihood of evidence for one model over another(Johnson et al., 2023). We intentionally chose not to utilize Bayes’ factor metrics because, in the authors’ opinions, they reduce inference to evaluation of a single metric over an arbitrary threshold, as opposed to analysis over posterior distributions of observed data. Furthermore, certain prior declarations yield undefined Bayes’ factors(Gelman and Rubin, 1995) potentially encouraging using suboptimum models in order to provide arbitrary decision metrics.

### Bayesian and Frequentist Approaches: A Holistic Approach to Inference

Following in the steps of Bayarri and Berger(Bayarri and Berger, 2004), data analysis should not consist solely of Bayesian or frequentist approaches devoid of the other. There are certainly cases where frequentist approaches should be used, such as clinical trials where preregistration and proper protocol design can provide bounds on false-positive and false negative rates necessary for translation of medical therapeutics. Hybrid frequentist and Bayesian approaches can also provide richer insight into analyses where posterior distributions are unidentifiable or difficult to sample(Raue et al., 2013) or in identifying when improper models have been chosen(Berger et al., 1997). Bayesian ideas of posterior predictive checks and model comparisons can also be applied to frequentist NHST, many of which would help address problems of replication and data transparency. As frequentist approaches are often baked into the pedagogy of neuroscience and neural engineering, we aim for this tutorial to be a thorough introduction into the application of Bayesian statistics to help develop a toolkit which can be used for robust data analysis or in conjunction with previously established frequentist approaches. These models are also easily extendable into Bayesian analogs of logistic or multinomial regressions, gaussian mixture models, Bayesian time series analyses, among many more.

### Code and Data Availability

The code/software and data described in this paper is freely available online at https://github.com/bscoventry/Practical-Bayesian-Inference-in-Neuroscience-Or-How-I-Learned-To-Stop-Worrying-and-Embrace-the-Dist. The code is available as Extended Data.

## Acknowledgements

The authors would like to thank the anonymous reviewers for their helpful and constructive feedback on this article. Funding for experiments which generated young and aged responses to SAM stimuli was provided by the National Institutes of Health (NIDCD R01DC011580, PI: ELB). Funding for experiments generating INS dose-response data was provided by the Purdue Institute for Integrative Neuroscience collaborative training grant (PI: BSC).

## Code and Data Repository

https://github.com/bscoventry/BayesianNeuralAnalysis

## Extended Data

All data and source code can be found in our GitHub toolbox: https://github.com/bscoventry/Practical-Bayesian-Inference-in-Neuroscience-Or-How-I-Learned-To-Stop-Worrying-and-Embrace-the-Dist.

Installation and Running of the Bayesian Inference Toolbox

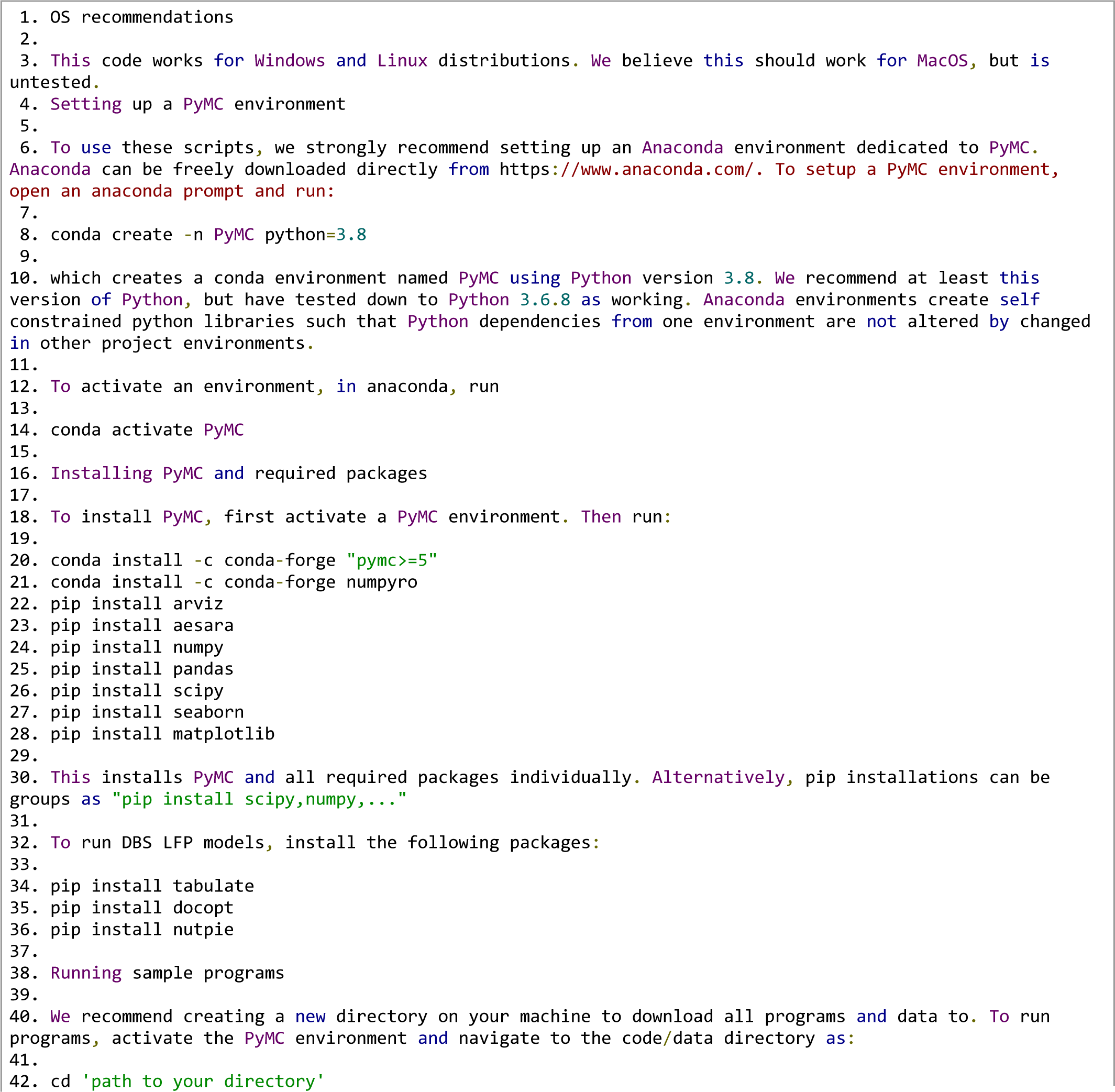
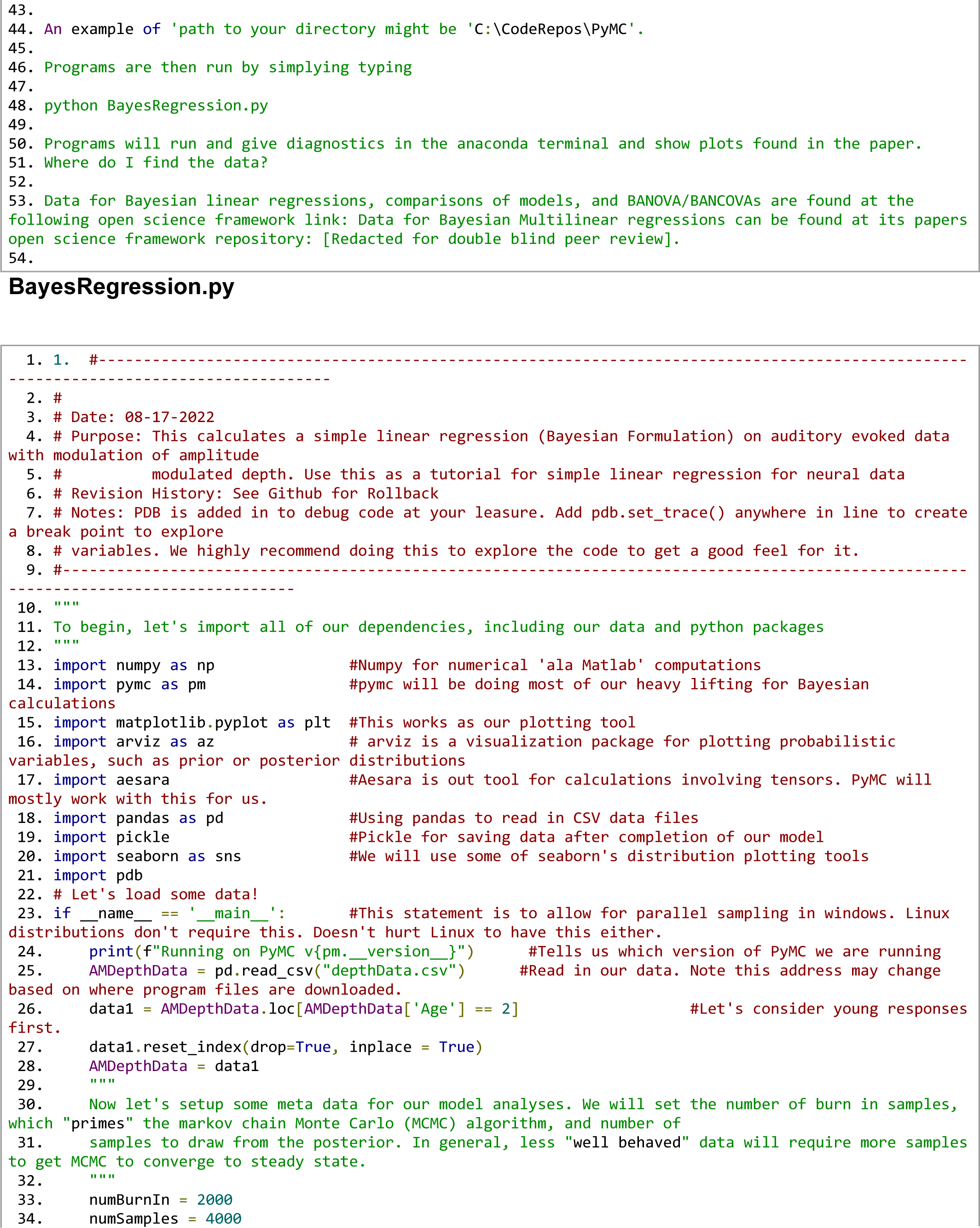
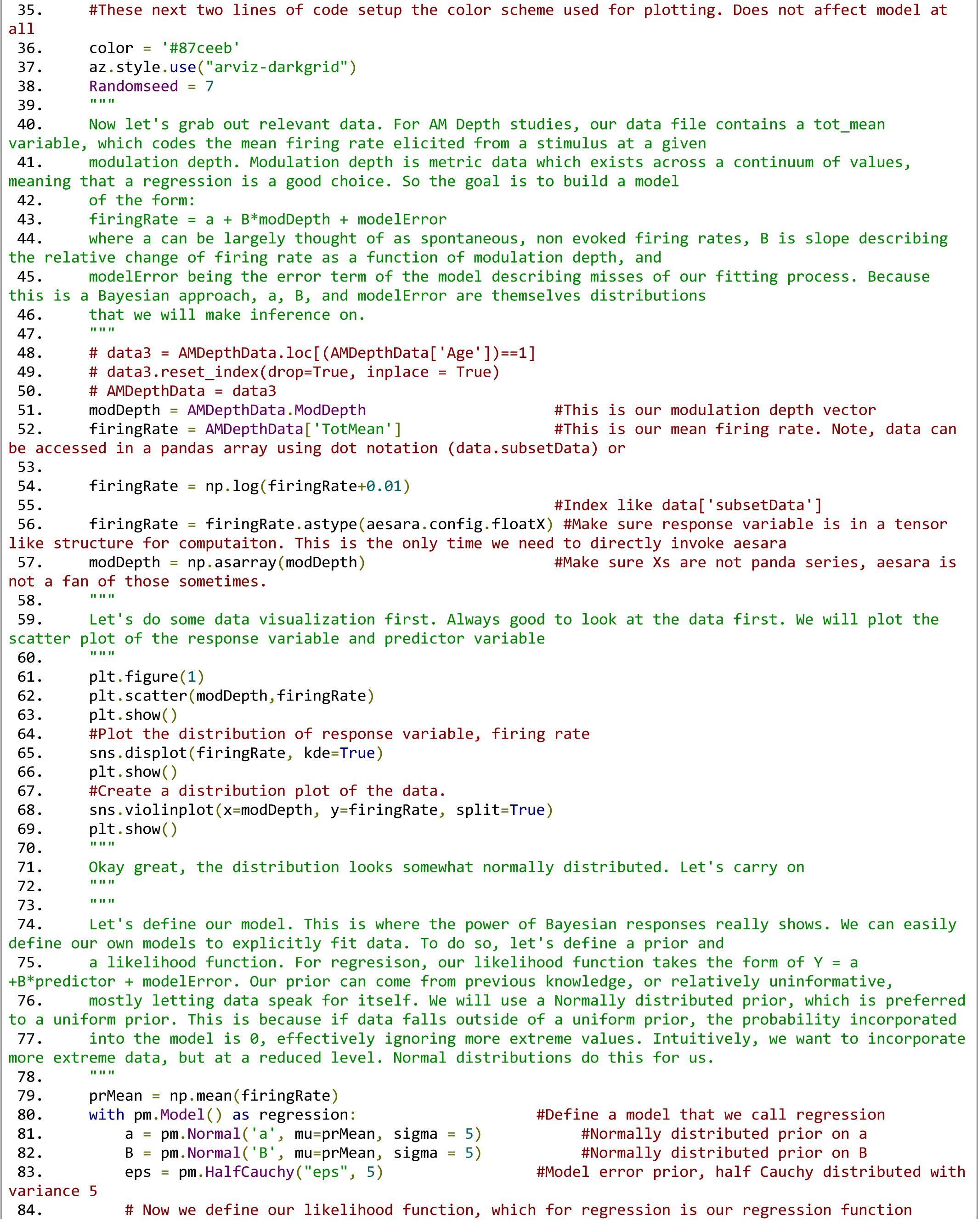
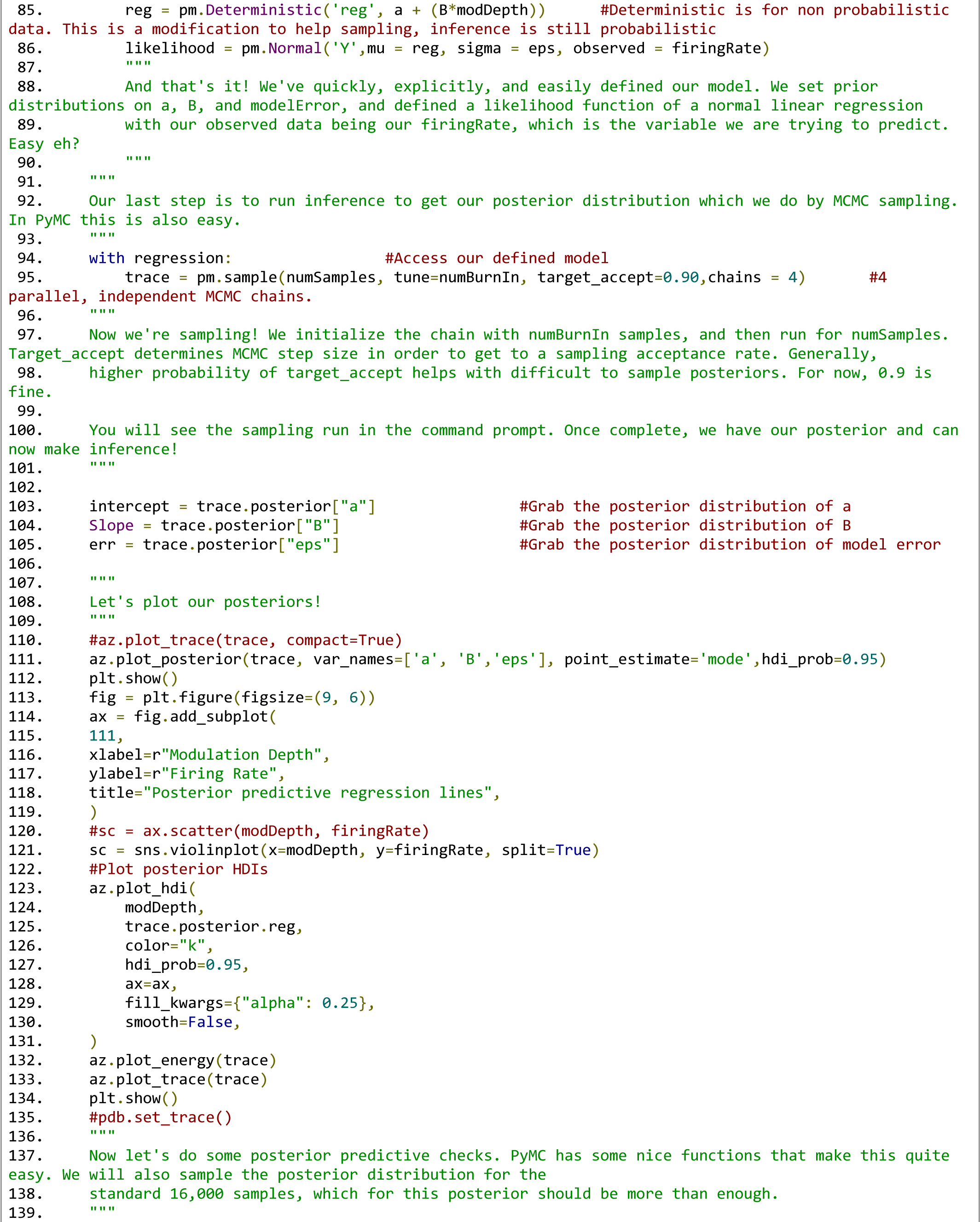
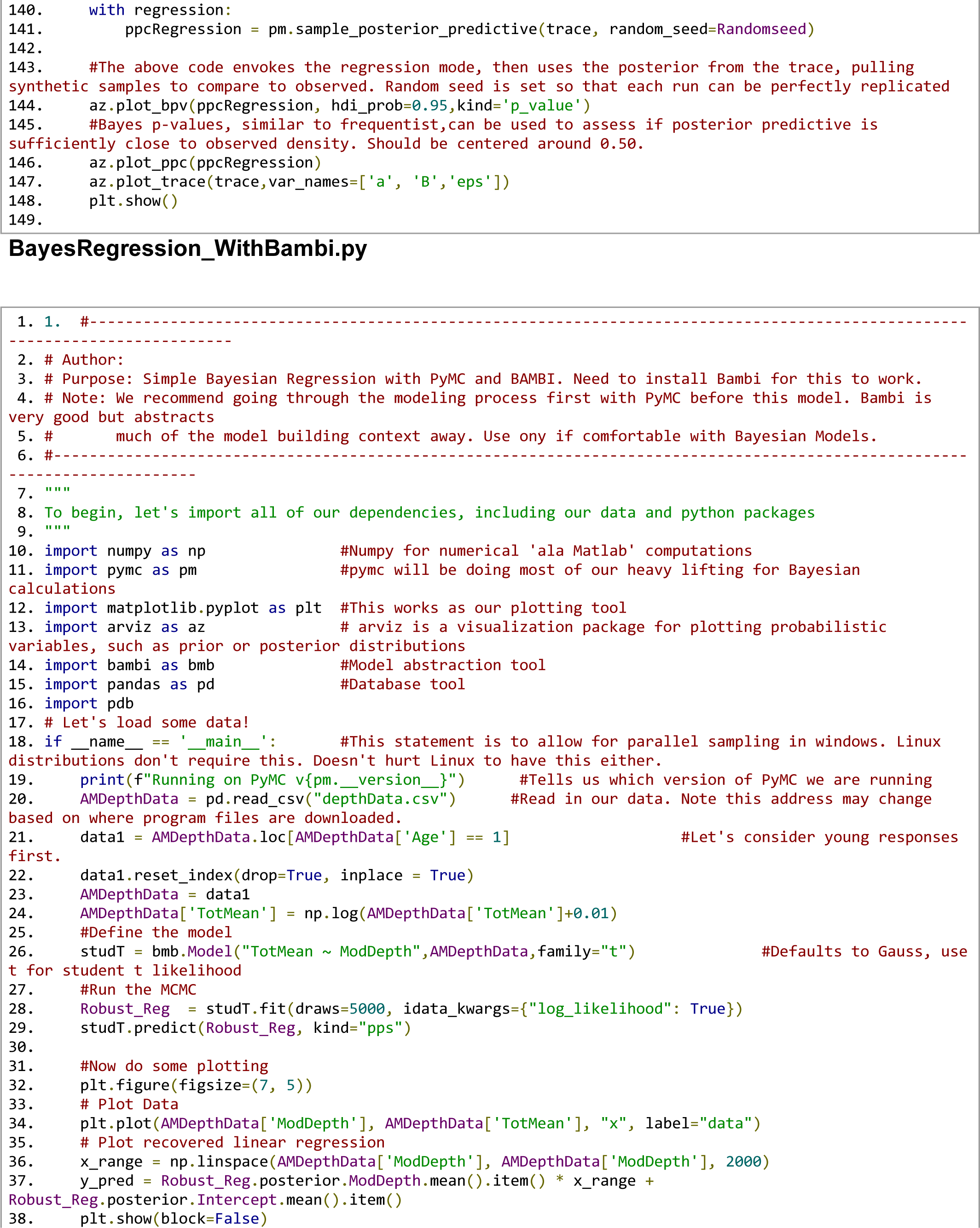
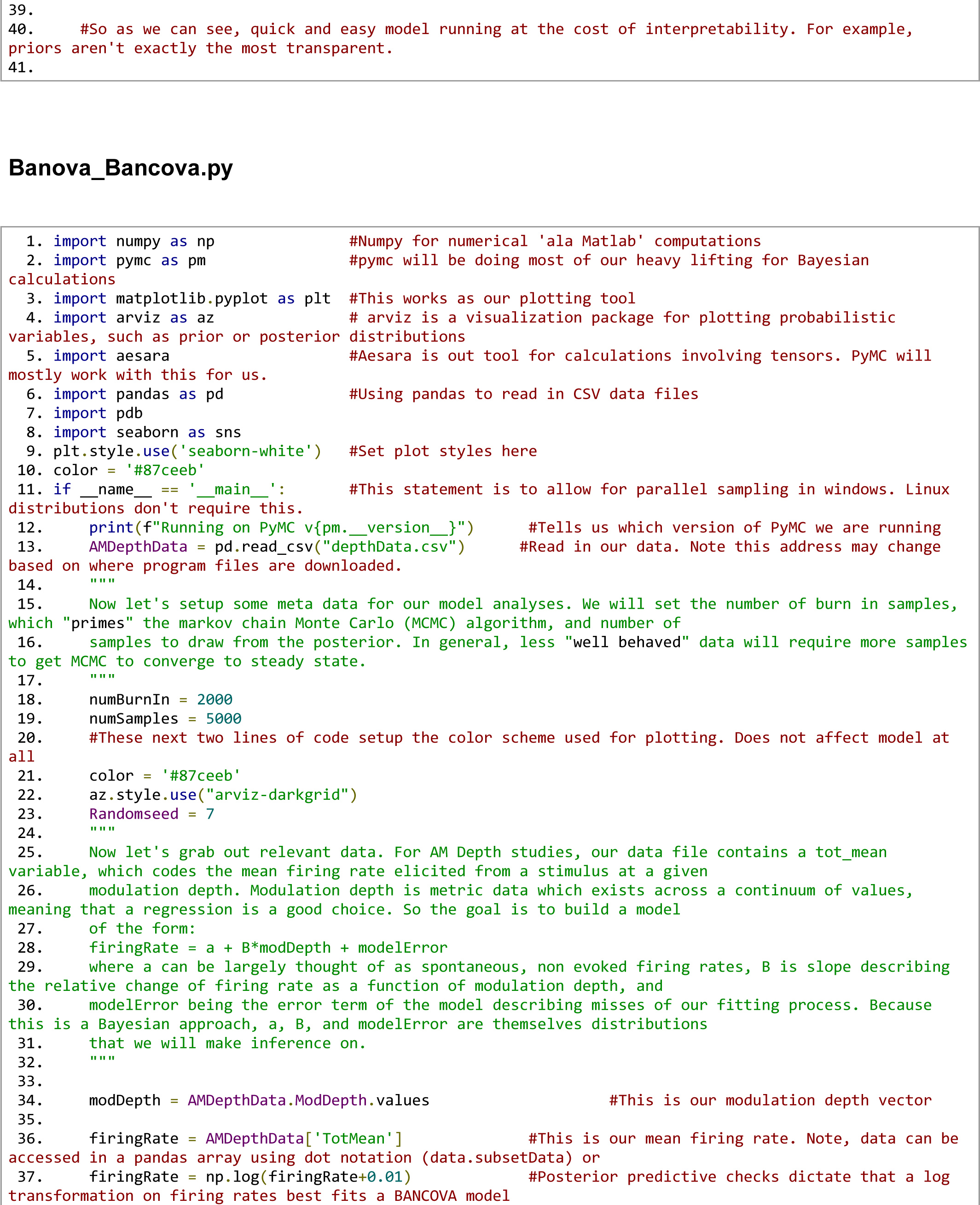
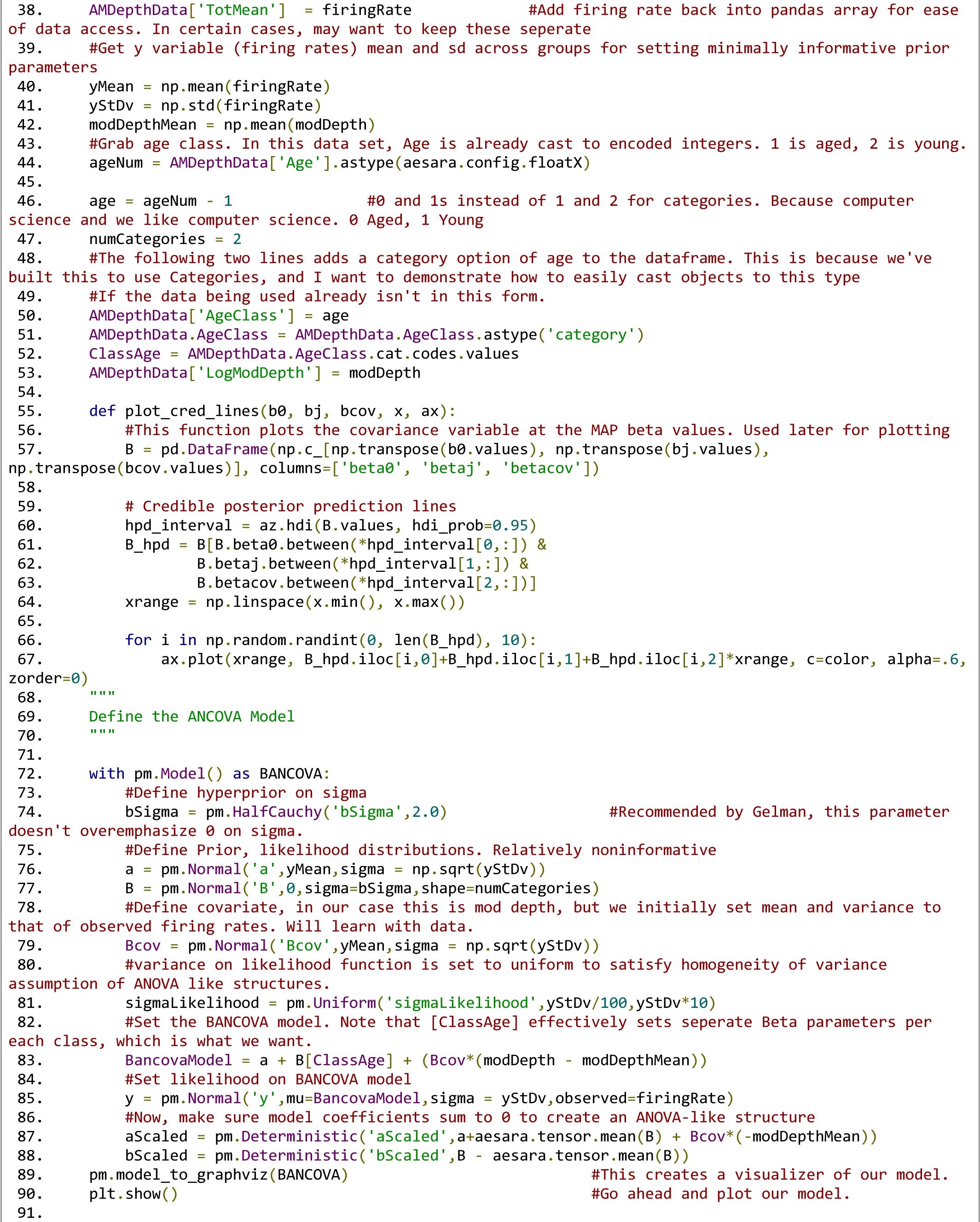
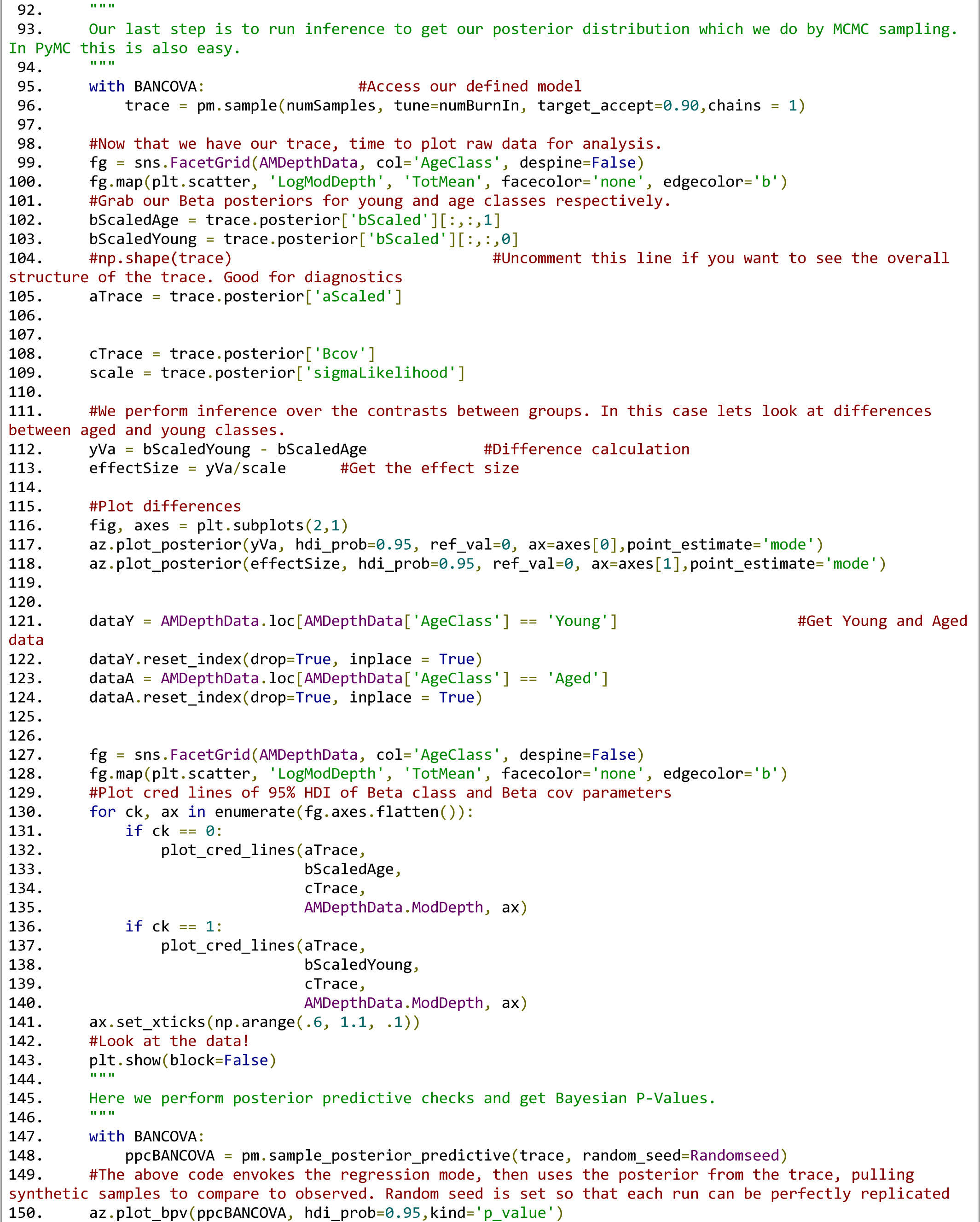
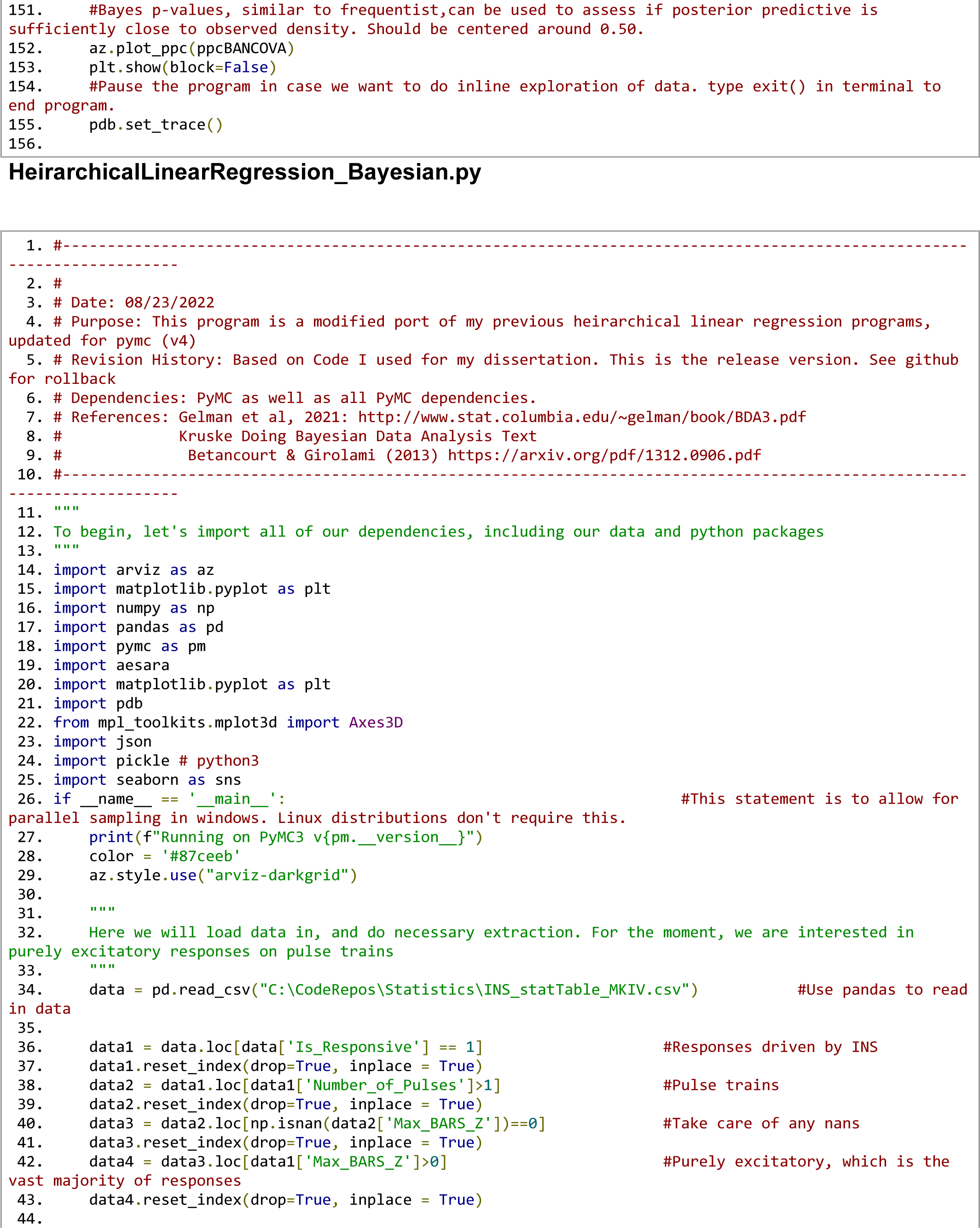
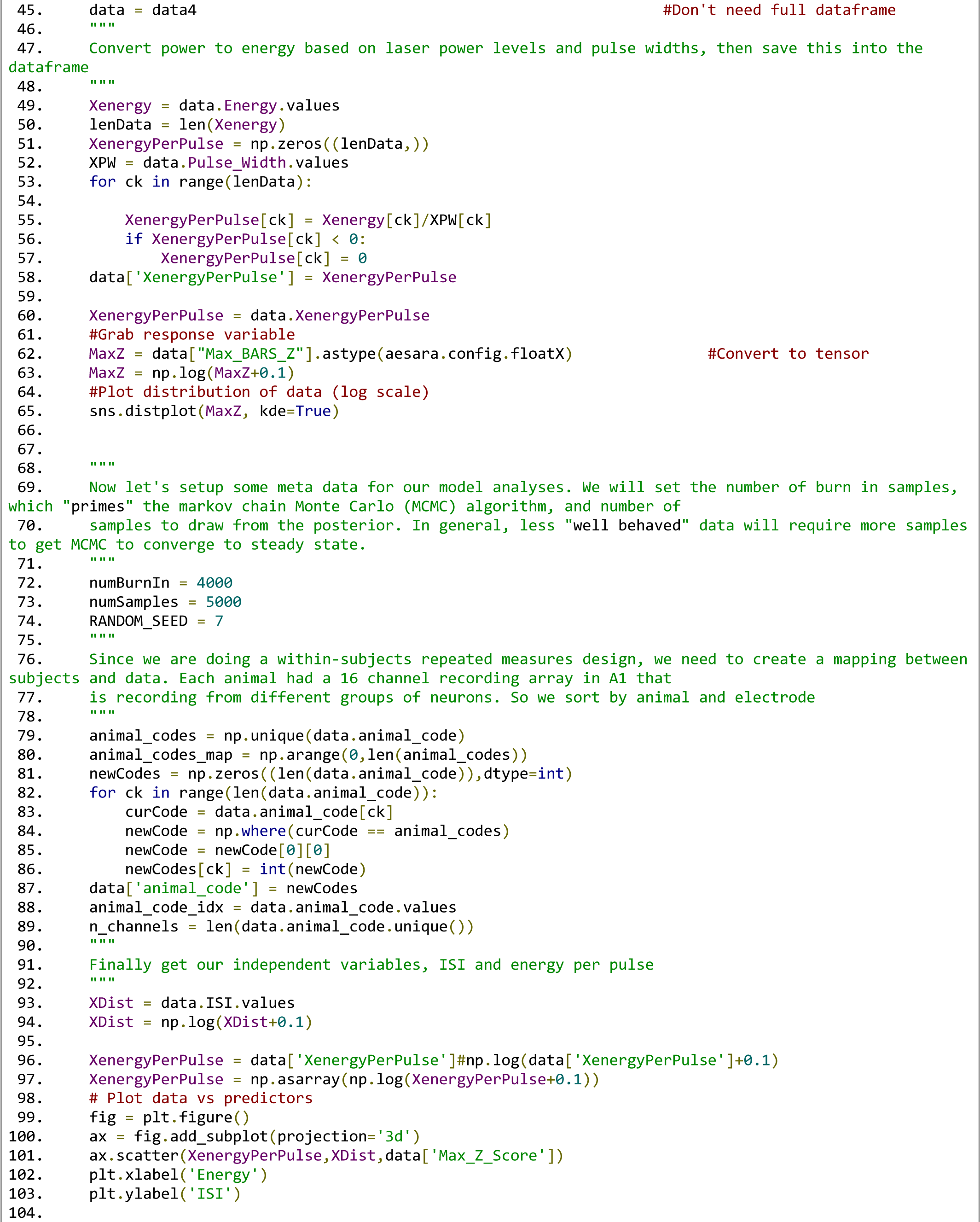
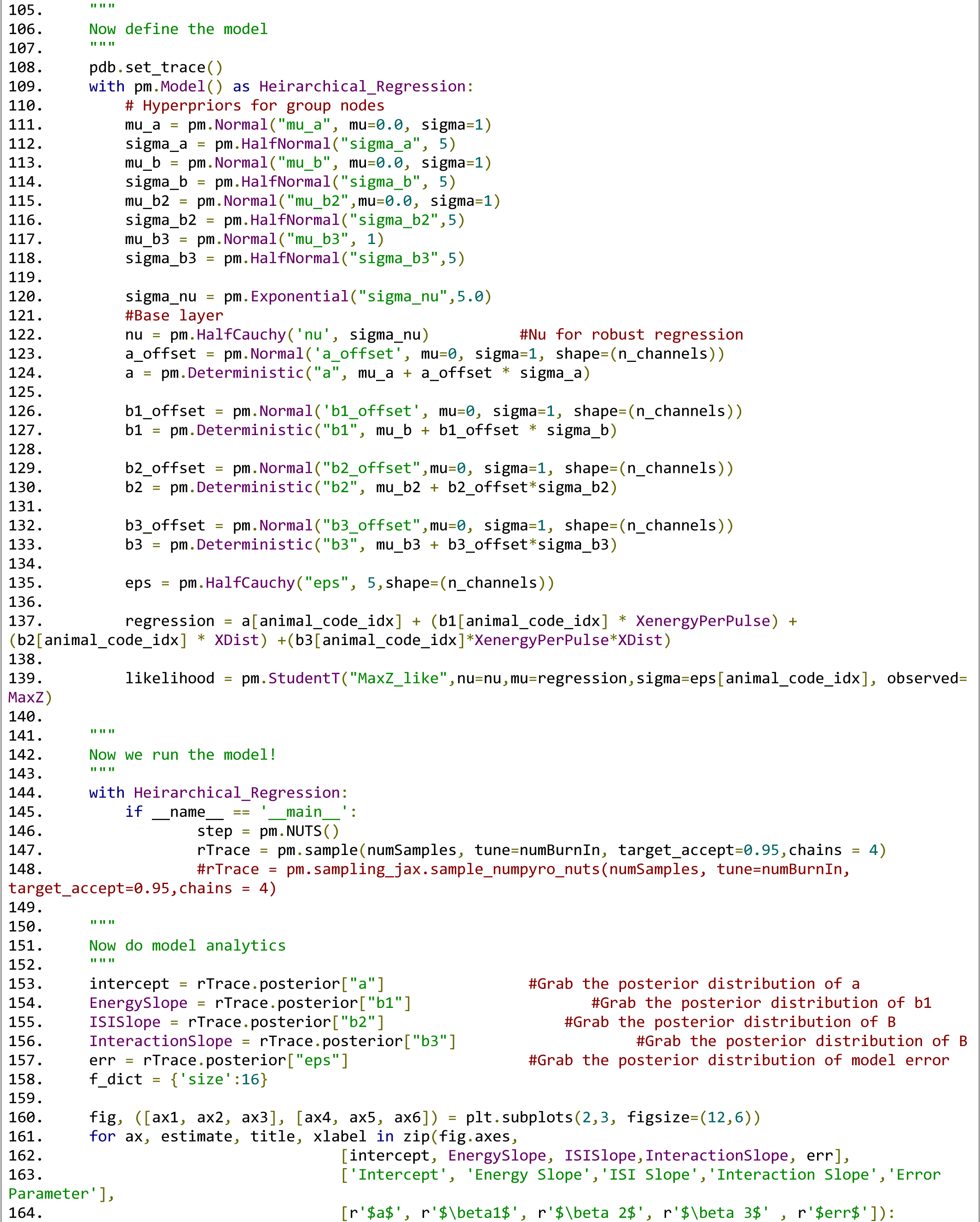
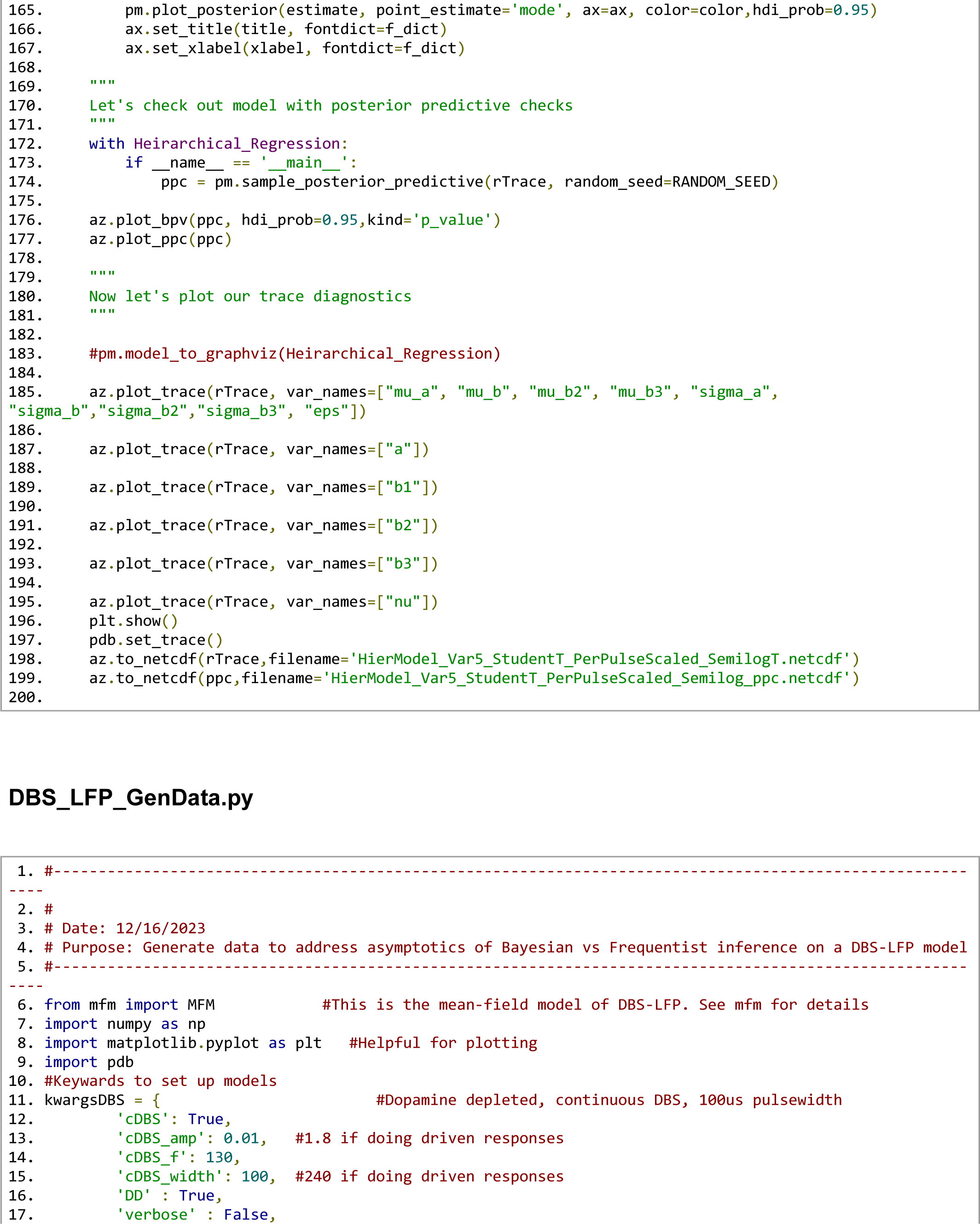
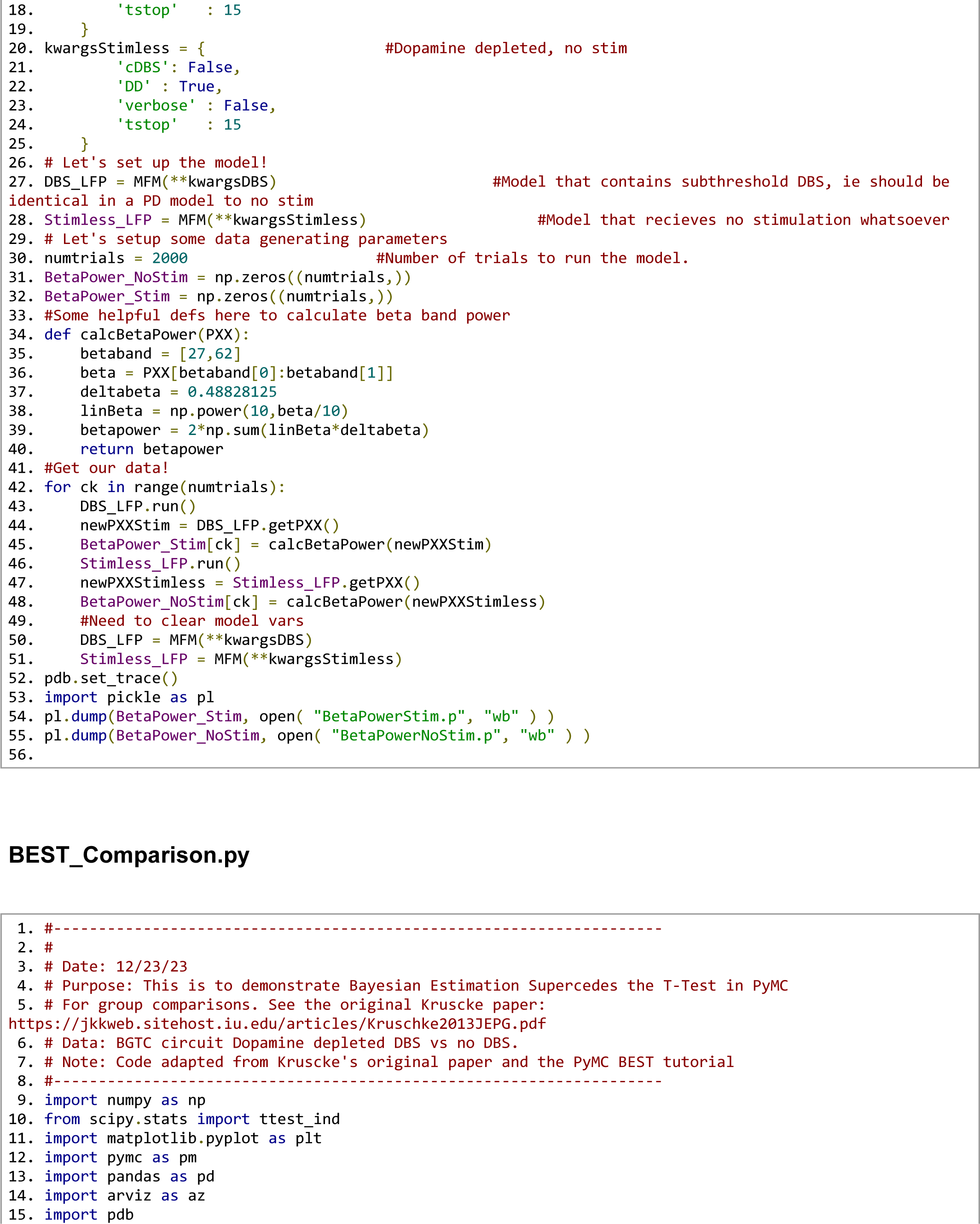
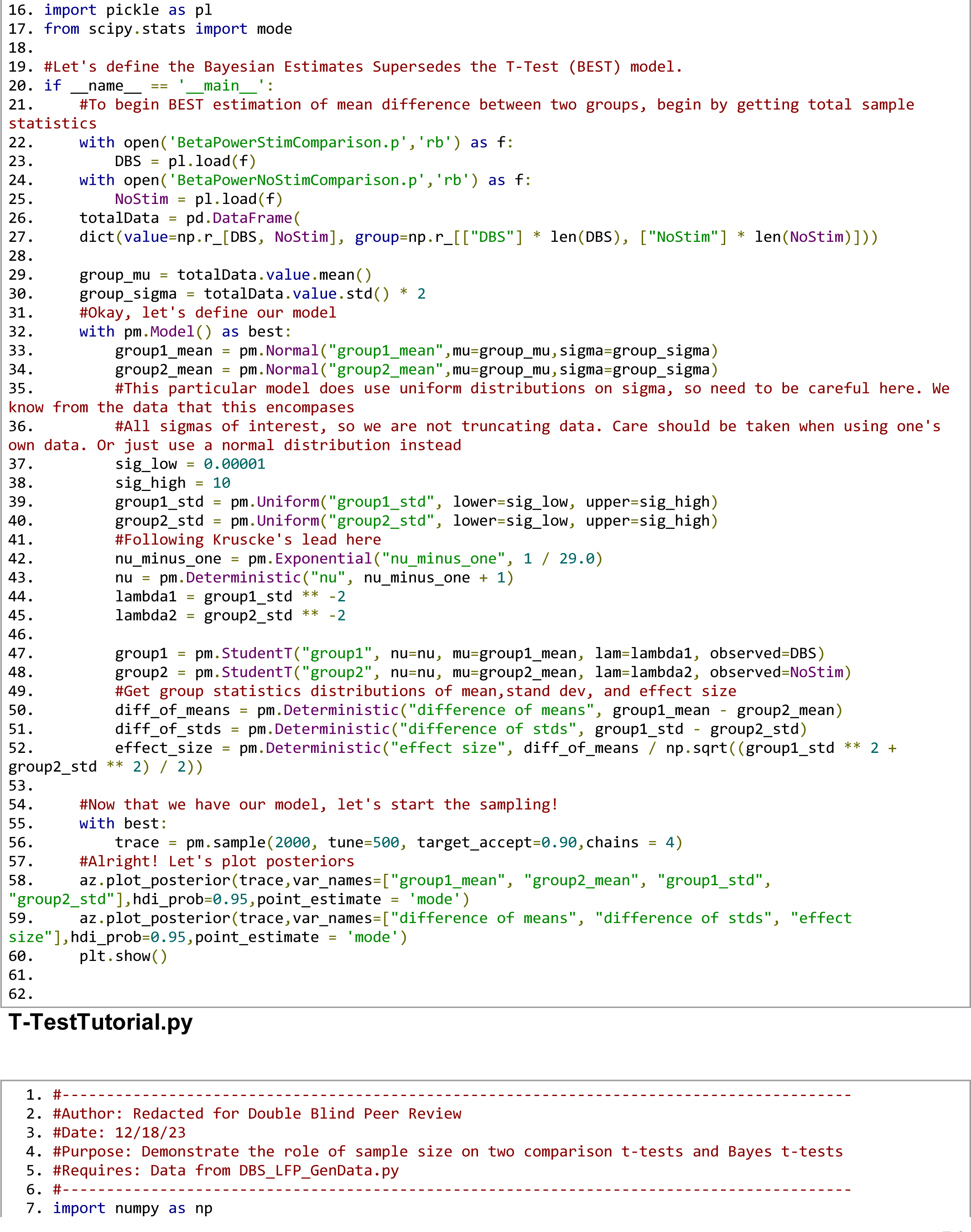
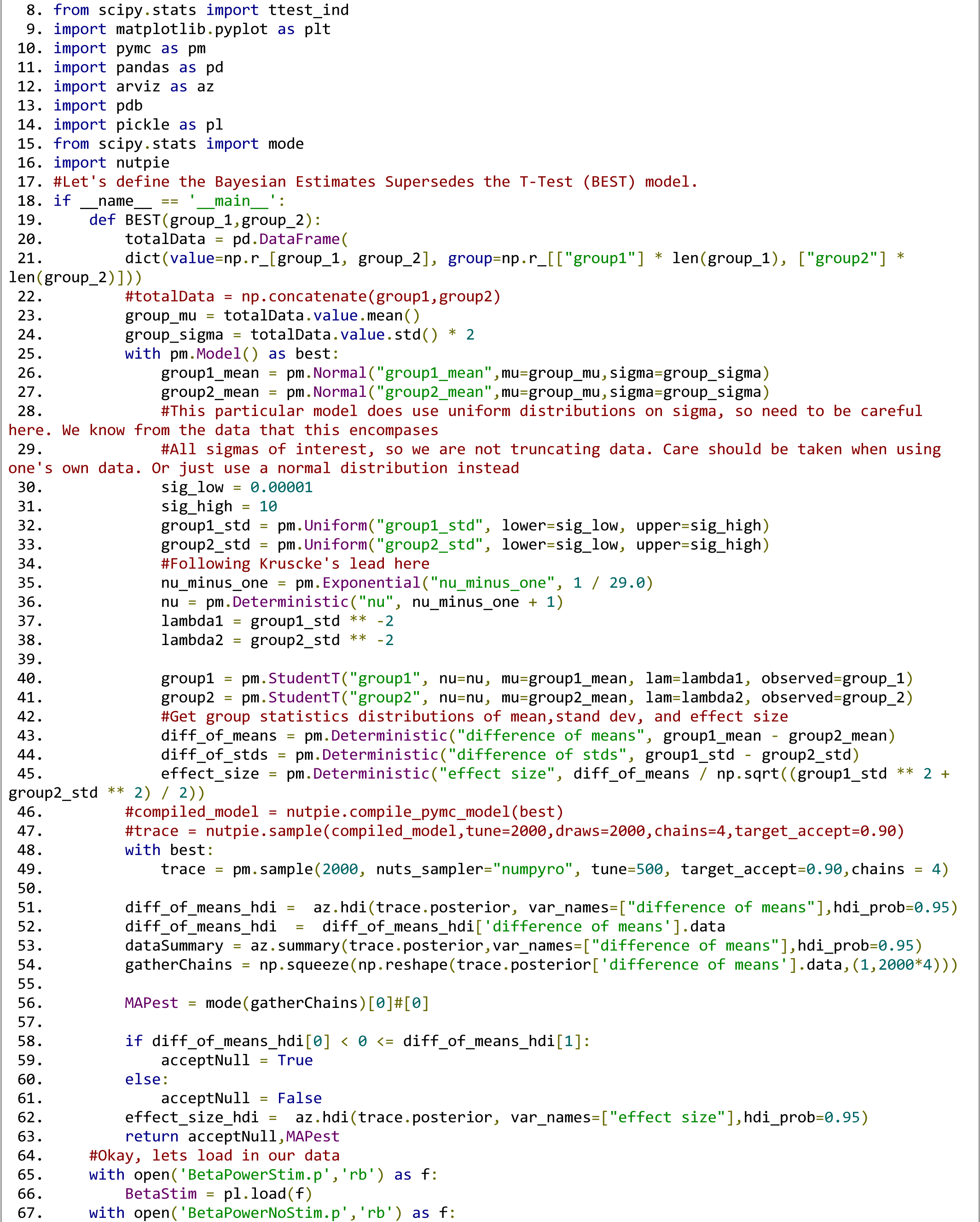
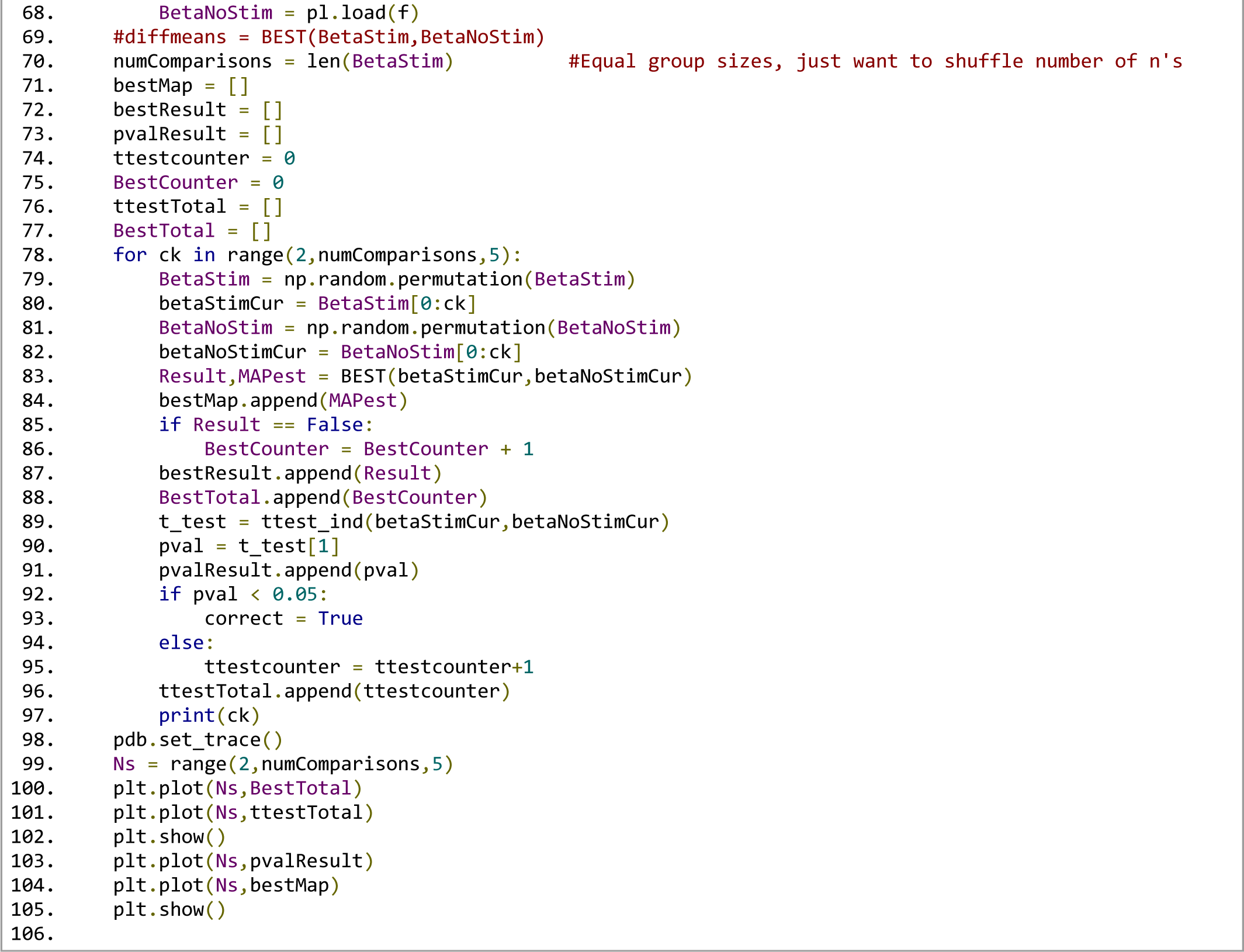

## References

Bartlett EL, Wang X (2007) Neural Representations of Temporally Modulated Signals in the Auditory Thalamus of Awake Primates. J Neurophysiol 97:1005–1017.

Bayarri MJ, Berger JO (2004) The Interplay of Bayesian and Frequentist Analysis. Stat Sci 19 Available at: https://projecteuclid.org/journals/statistical-science/volume-19/issue-1/The-Interplay-of-Bayesian-and-Frequentist-Analysis/10.1214/088342304000000116.full [Accessed August 5, 2023].

Berger JO, Boukai B, Wang Y (1997) Unified frequentist and Bayesian testing of a precise hypothesis. Stat Sci 12 Available at: https://projecteuclid.org/journals/statistical-science/volume-12/issue-3/Unified-frequentist-and-Bayesian-testing-of-a-precise-hypothesis/10.1214/ss/1030037904.full [Accessed August 8, 2023].

Betancourt M (2017) A Conceptual Introduction to Hamiltonian Monte Carlo. Available at: https://arxiv.org/abs/1701.02434 [Accessed July 26, 2023].

Bielza C, Larranaga P (2014) Bayesian networks in neuroscience: a survey. Front Comput Neurosci 8 Available at: http://journal.frontiersin.org/article/10.3389/fncom.2014.00131/abstract [Accessed November 2, 2023].

Bishop C (2006) Pattern Recognition and Machine Learning, 1st ed. New York, NY: Springer.

Blackwell DL, Ramamoorthi RV (1982) A Bayes but Not Classically Sufficient Statistic. Ann Stat 10:1025–1026.

Bove C, Travagli RA (2019) Neurophysiology of the brain stem in Parkinson’s disease. J Neurophysiol 121:1856–1864.

GEP, Tiao GC (2011) Bayesian Inference in Statistical Analysis, 1st ed. Germany: Wiley.

Brooks SP (2003) Bayesian Computation: A Statistical Revolution. Philos Trans Math Phys Eng Sci 361:2681–2697.

Button KS, Ioannidis JPA, Mokrysz C, Nosek BA, Flint J, Robinson ESJ, Munafò MR (2013) Power failure: why small sample size undermines the reliability of neuroscience. Nat Rev Neurosci 14:365–376.

Bzdok D, Ioannidis JPA (2019) Exploration, Inference, and Prediction in Neuroscience and Biomedicine. Trends Neurosci 42:251–262.

Cant NB, Benson CG (2006) Organization of the inferior colliculus of the gerbil (Meriones unguiculatus): Differences in distribution of projections from the cochlear nuclei and the superior olivary complex. J Comp Neurol 495:511–528.

Capretto T, Piho C, Kumar R, Westfall J, Yarkoni T, Martin OA (2022) **Bambi** : A Simple Interface for Fitting Bayesian Linear Models in Python. J Stat Softw 103 Available at: https://www.jstatsoft.org/v103/i15/ [Accessed August 29, 2023].

Carpenter B, Gelman A, Hoffman MD, Lee D, Goodrich B, Betancourt M, Brubaker M, Guo J, Li P, Riddell A (2017) Stan : A Probabilistic Programming Language. J Stat Softw 76 Available at: http://www.jstatsoft.org/v76/i01/ [Accessed August 5, 2023].

Caspary DM, Palombi PS, Hughes LF (2002) GABAergic inputs shape responses to amplitude modulated stimuli in the inferior colliculus. Hear Res 168:163–173.

Cayce JM, Friedman RM, Chen G, Jansen ED, Mahadevan-Jansen A, Roe AW (2014) Infrared neural stimulation of primary visual cortex in non-human primates. NeuroImage 84:181–190.

Cayce JM, Friedman RM, Jansen ED, Mahavaden-Jansen A, Roe AW (2011) Pulsed infrared light alters neural activity in rat somatosensory cortex in vivo. NeuroImage 57:155–166.

Cinotti F, Humphries MD (2022) Bayesian Mapping of the Striatal Microcircuit Reveals Robust Asymmetries in the Probabilities and Distances of Connections. J Neurosci 42:1417–1435.

Colombo M, Seriès P (2012) Bayes in the Brain—On Bayesian Modelling in Neuroscience. Br J Philos Sci 63:697–723.

Coventry BS, Bartlett EL (2023) Closed-Loop Reinforcement Learning Based Deep Brain Stimulation Using SpikerNet: A Computational Model. In: 11th International IEEE EMBS Conference on Neural Engineering, pp 1–4. Baltimore, Maryland USA.

Coventry BS, Lawlor GL, Bagnati CB, Krogmeier C, Bartlett EL (2024) Characterization and closed-loop control of infrared thalamocortical stimulation produces spatially constrained single-unit responses. PNAS 3(2) Nexus:pgae082.

Coventry BS, Sick JT, Talavage TM, Stantz KM, Bartlett EL (2020) Short-wave Infrared Neural Stimulation Drives Graded Sciatic Nerve Activation Across A Continuum of Wavelengths. In: 2020 42nd Annual International Conference of the IEEE Engineering in Medicine & Biology Society (EMBC). IEEE.

Cronin B, Stevenson IH, Sur M, Körding KP (2010) Hierarchical Bayesian Modeling and Markov Chain Monte Carlo Sampling for Tuning-Curve Analysis. J Neurophysiol 103:591–602.

De La Rocha J, Doiron B, Shea-Brown E, Josić K, Reyes A (2007) Correlation between neural spike trains increases with firing rate. Nature 448:802–806.

Dickson DW (2018) Neuropathology of Parkinson disease. Parkinsonism Relat Disord 46:S30–S33.

Etz A (2018) Introduction to the Concept of Likelihood and Its Applications. Adv Methods Pract Psychol Sci 1:60–69.

Falowski SM, Sharan A, Reyes BAS, Sikkema C, Szot P, Van Bockstaele EJ (2011) An Evaluation of Neuroplasticity and Behavior After Deep Brain Stimulation of the Nucleus Accumbens in an Animal Model of Depression. Neurosurgery 69:1281–1290.

Fienberg SE (2006) When did Bayesian inference become “Bayesian”? Bayesian Anal 1 Available at: https://projecteuclid.org/journals/bayesian-analysis/volume-1/issue-1/When-did-Bayesian-inference-become-Bayesian/10.1214/06-BA101.full [Accessed October 23, 2023].

Fisher RA (1992) Statistical Methods for Research Workers. In: Breakthroughs in Statistics (Kotz S, Johnson NL, eds), pp 66–70 Springer Series in Statistics. New York, NY: Springer New York. Available at: http://link.springer.com/10.1007/978-1-4612-4380-9_6 [Accessed August 5, 2023].

Frisina DR, Frisina RD (1997) Speech recognition in noise and presbycusis: relations to possible neural mechanisms. Hear Res 106:95–104.

Gelman A (2005) Analysis of variance—why it is more important than ever. Ann Stat 33 Available at: https://projecteuclid.org/journals/annals-of-statistics/volume-33/issue-1/Analysis-of-variancewhy-it-is-more-important-than-ever/10.1214/009053604000001048.full [Accessed August 6, 2023].

Gelman A (2006) Prior distributions for variance parameters in hierarchical models (comment on article by Browne and Draper). Bayesian Anal 1 Available at: https://projecteuclid.org/journals/bayesian-analysis/volume-1/issue-3/Prior-distributions-for-variance-parameters-in-hierarchical-models-comment-on/10.1214/06-BA117A.full.

Gelman A, Carlin J, Stern H, Dunson D, Vehtari A, Rubin D (2021) Bayesian Data Analysis, 3rd ed. Boca Raton: Chapman and Hall/CRC.

Gelman A, Hill J, Yajima M (2009) Why we (usually) don’t have to worry about multiple comparisons. Available at: https://arxiv.org/abs/0907.2478 [Accessed August 29, 2023].

Gelman A, Hwang J, Vehtari A (2014) Understanding predictive information criteria for Bayesian models. Stat Comput 24:997–1016.

Gelman A, Rubin DB (1992) Inference from Iterative Simulation Using Multiple Sequences. Stat Sci 7:457–472.

Gelman A, Rubin DB (1995) Avoiding Model Selection in Bayesian Social Research. Sociol Methodol 25:165.

Gelman A, Shalizi CR (2013) Philosophy and the practice of Bayesian statistics: *Philosophy and the practice of Bayesian statistics*. Br J Math Stat Psychol 66:8–38.

Gerwinn S, Macke JH, Bethge M (2010) Bayesian inference for generalized linear models for spiking neurons. Front Comput Neurosci 4 Available at: http://journal.frontiersin.org/article/10.3389/fncom.2010.00012/abstract [Accessed November 2, 2023].

Gilks WR, Richardson S, Spiegelhalter DJ (1996) Markov Chain Monte Carlo in Practice. Boca Raton: Chapman and Hall/CRC.

Grado LL, Johnson MD, Netoff TI (2018) Bayesian adaptive dual control of deep brain stimulation in a computational model of Parkinson’s disease Santaniello S, ed. PLOS Comput Biol 14:e1006606.

Gregory BA, Thompson CH, Salatino JW, Railing MJ, Zimmerman AF, Gupta B, Williams K, Beatty JA, Cox CL, Purcell EK (2022) Structural and functional changes of pyramidal neurons at the site of an implanted microelectrode array in rat primary motor cortex. Available at: http://biorxiv.org/lookup/doi/10.1101/2022.09.15.507997 [Accessed March 31, 2023].

Grimsley CA, Sanchez JT, Sivaramakrishnan S (2013) Midbrain local circuits shape sound intensity codes. Front Neural Circuits 7 Available at: http://journal.frontiersin.org/article/10.3389/fncir.2013.00174/abstract [Accessed August 27, 2023].

Guidetti M, Marceglia S, Loh A, Harmsen IE, Meoni S, Foffani G, Lozano AM, Moro E, Volkmann J, Priori A (2021) Clinical perspectives of adaptive deep brain stimulation. Brain Stimulat 14:1238–1247.

Herrmann B, Parthasarathy A, Bartlett EL (2017) Ageing affects dual encoding of periodicity and envelope shape in rat inferior colliculus neurons Foxe J, ed. Eur J Neurosci 45:299–311.

Hoffman MD, Gelman A (2011) The No-U-Turn Sampler: Adaptively Setting Path Lengths in Hamiltonian Monte Carlo. Available at: https://arxiv.org/abs/1111.4246 [Accessed July 26, 2023].

Izzo AD, Suh E, Pathria J, Walsh JT, Whitlon DS, Richter C-P (2007) Selectivity of neural stimulation in the auditory system: a comparison of optic and electric stimuli. J Biomed Opt 12:021008.

Johnson VE, Pramanik S, Shudde R (2023) Bayes factor functions for reporting outcomes of hypothesis tests. Proc Natl Acad Sci 120:e2217331120.

Kelly JB, Caspary DM (2005) Pharmacology of the inferior colliculus. In: The inferior colliculus., 1st ed. New York, NY: Springer.

Krueger JI, Heck PR (2019) Putting the *P* -Value in its Place. Am Stat 73:122–128.

Kruschke JK (2010) What to believe: Bayesian methods for data analysis. Trends Cogn Sci 14:293–300.

Kruschke JK (2011) Bayesian Assessment of Null Values Via Parameter Estimation and Model Comparison. Perspect Psychol Sci 6:299–312.

Kruschke JK (2013) Bayesian estimation supersedes the t test. J Exp Psychol Gen 142:573–603.

Kruschke JK (2014) Doing Bayesian Data Analysis: A tutorial with R, JAGS, and stan, 2nd ed. Academic Press. Available at: http://www.indiana.edu/~kruschke/DoingBayesianDataAnalysis/.

Kruschke JK (2018) Rejecting or Accepting Parameter Values in Bayesian Estimation. Adv Methods Pract Psychol Sci 1:270–280.

Kruschke JK (2021) Bayesian Analysis Reporting Guidelines. Nat Hum Behav 5:1282–1291.

Kruschke JK, Vanpaemel W (2015) Bayesian Estimation in Hierarchical Models. In: The Oxford Handbook of Computational and Mathematical Psychology, pp 279–299. Oxford University Press.

Kutner MH, Nachtsheim CJ, Neter J, Li W (2005) Applied Linear Statistical Models, 5th ed. Boston, MA: McGraw-Hill Irwin.

Little S, Pogosyan A, Neal S, Zavala B, Zrinzo L, Hariz M, Foltynie T, Limousin P, Ashkan K, FitzGerald J, Green AL, Aziz TZ, Brown P (2013) Adaptive deep brain stimulation in advanced Parkinson disease. Ann Neurol 74:449–457.

Loftus WC, Bishop DC, Oliver DL (2010) Differential Patterns of Inputs Create Functional Zones in Central Nucleus of Inferior Colliculus. J Neurosci 30:13396–13408.

Love J, Selker R, Marsman M, Jamil T, Dropmann D, Verhagen J, Ly A, Gronau QF, Smíra M, Epskamp S, Matzke D, Wild A, Knight P, Rouder JN, Morey RD, Wagenmakers E-J (2019) **JASP** : Graphical Statistical Software for Common Statistical Designs. J Stat Softw 88 Available at: http://www.jstatsoft.org/v88/i02/ [Accessed August 5, 2023].

Ma WJ (2019) Bayesian Decision Models: A Primer. Neuron 104:164–175.

Maris E (2012) Statistical testing in electrophysiological studies. Psychophysiology 49:549–565.

Munafò MR, Nosek BA, Bishop DVM, Button KS, Chambers CD, Percie Du Sert N, Simonsohn U, Wagenmakers E-J, Ware JJ, Ioannidis JPA (2017) A manifesto for reproducible science. Nat Hum Behav 1:0021.

Nieuwenhuis S, Forstmann BU, Wagenmakers E-J (2011) Erroneous analyses of interactions in neuroscience: a problem of significance. Nat Neurosci 14:1105–1107.

Nuzzo R (2014) Scientific method: Statistical errors. Nature 506:150–152.

Palombi PS, Backoff PM, Caspary DM (2001) Responses of young and aged rat inferior colliculus neurons to sinusoidally amplitude modulated stimuli. Hear Res 153:174–180.

Paninski L, Fellows MR, Hatsopoulos NG, Donoghue JP (2004) Spatiotemporal Tuning of Motor Cortical Neurons for Hand Position and Velocity. J Neurophysiol 91:515–532.

Park M, Pillow JW (2011) Receptive Field Inference with Localized Priors Sporns O, ed. PLoS Comput Biol 7:e1002219.

Parthasarathy A, Bartlett E (2012) Two-channel recording of auditory-evoked potentials to detect age-related deficits in temporal processing. Hear Res 289:52–62.

Parthasarathy A, Cunningham PA, Bartlett EL (2010) Age-Related Differences in Auditory Processing as Assessed by Amplitude-Modulation Following Responses in Quiet and in Noise. Front Aging Neurosci 2 Available at: http://journal.frontiersin.org/article/10.3389/fnagi.2010.00152/abstract [Accessed March 13, 2023].

Parzen E (1962) On Estimation of a Probability Density Function and Mode. Ann Math Stat 33:1065– 1076.

Polson NG, Scott JG (2012) On the Half-Cauchy Prior for a Global Scale Parameter. Bayesian Anal 7 Available at: https://projecteuclid.org/journals/bayesian-analysis/volume-7/issue-4/On-the-Half-Cauchy-Prior-for-a-Global-Scale-Parameter/10.1214/12-BA730.full [Accessed August 29, 2023].

Quiroga RQ, Nadasdy Z, Ben-Shaul Y (2004) Unsupervised spike detection and sorting with wavelets and superparamagnetic clustering. Neural Comput 16:1661–1687.

Rabang CF, Parthasarathy A, Venkataraman Y, Fisher ZL, Gardner SM, Bartlett EL (2012) A computational model of inferior colliculus responses to amplitude modulated sounds in young and aged rats. Front Neural Circuits 6:77.

Raue A, Kreutz C, Theis FJ, Timmer J (2013) Joining forces of Bayesian and frequentist methodology: a study for inference in the presence of non-identifiability. Philos Trans R Soc Math Phys Eng Sci 371:20110544.

Rosenblatt M (1956) Remarks on Some Nonparametric Estimates of a Density Function. Ann Math Stat 27:832–837.

Salvatier J, Wiecki TV, Fonnesbeck C (2016) Probabilistic programming in Python using PyMC3. PeerJ Comput Sci 2:e55.

Simon H, Frisina RD, Walton JP (2004) Age reduces response latency of mouse inferior colliculus neurons to AM sounds. J Acoust Soc Am 116:469–477.

Singh A (2018) Oscillatory activity in the cortico-basal ganglia-thalamic neural circuits in Parkinson’s disease. Eur J Neurosci 48:2869–2878.

Smid SC, McNeish D, Miočević M, Van De Schoot R (2020) Bayesian Versus Frequentist Estimation for Structural Equation Models in Small Sample Contexts: A Systematic Review. Struct Equ Model Multidiscip J 27:131–161.

Song S, Regan B, Ereifej ES, Chan ER, Capadona JR (2022) Neuroinflammatory Gene Expression Analysis Reveals Pathways of Interest as Potential Targets to Improve the Recording Performance of Intracortical Microelectrodes. Cells 11:2348.

Van Albada SJ, Robinson PA (2009) Mean-field modeling of the basal ganglia-thalamocortical system. I. J Theor Biol 257:642–663.

Van De Schoot R, Depaoli S, King R, Kramer B, Märtens K, Tadesse MG, Vannucci M, Gelman A, Veen D, Willemsen J, Yau C (2021) Bayesian statistics and modelling. Nat Rev Methods Primer 1:1.

Van Kuyck K, Welkenhuysen M, Arckens L, Sciot R, Nuttin B (2007) Histological Alterations Induced by Electrode Implantation and Electrical Stimulation in the Human Brain: A Review. Neuromodulation Technol Neural Interface 10:244–261.

Vasquez-Lopez SA, Weissenberger Y, Lohse M, Keating P, King AJ, Dahmen JC (2017) Thalamic input to auditory cortex is locally heterogeneous but globally tonotopic. eLife 6:e25141.

Vehtari A, Gelman A, Gabry J (2017) Practical Bayesian model evaluation using leave-one-out cross-validation and WAIC. Stat Comput 27:1413–1432.

Vogelstein JT, Watson BO, Packer AM, Yuste R, Jedynak B, Paninski L (2009) Spike Inference from Calcium Imaging Using Sequential Monte Carlo Methods. Biophys J 97:636–655.

Wagenmakers E-J (2007) A practical solution to the pervasive problems of p values. Psychon Bull Rev 14:779–804.

Wells J, Kao C, Jansen ED, Konrad P, Mahadevan-Jansen A (2005) Application of infrared light for in vivo neural stimulation. J Biomed Opt 10:064003.

Wood E, Fellows M, Donoghue JR, Black MJ (2004) Automatic spike sorting for neural decoding. In: The 26th Annual International Conference of the IEEE Engineering in Medicine and Biology Society, pp 4009–4012. San Francisco, CA, USA: IEEE. Available at: http://ieeexplore.ieee.org/document/1404120/ [Accessed January 31, 2024].

Woolley AJ, Desai HA, Otto KJ (2013) Chronic intracortical microelectrode arrays induce non-uniform, depth-related tissue responses. J Neural Eng 10:026007.

Wu W, Gao Y, Bienenstock E, Donoghue JP, Black MJ (2006) Bayesian Population Decoding of Motor Cortical Activity Using a Kalman Filter. Neural Comput 18:80–118.

